# Limited evidence for probabilistic cueing effects on grating-evoked event-related potentials and orientation decoding performance

**DOI:** 10.1101/2024.05.26.595980

**Authors:** Carla den Ouden, Máire Kashyap, Morgan Kikkawa, Daniel Feuerriegel

**Affiliations:** Melbourne School of Psychological Sciences, The University of Melbourne

**Keywords:** EEG, ERP, expectation, MVPA, surprise

## Abstract

We can rapidly learn recurring patterns that occur within our sensory environments. This knowledge allows us to form expectations about future sensory events. Several influential predictive coding models posit that, when a stimulus matches our expectations, the activity of feature-selective neurons in visual cortex will be suppressed relative to when that stimulus is unexpected. However, after accounting for known critical confounds, there is currently scant evidence for these hypothesised effects from studies recording electrophysiological neural activity. To provide a strong test for expectation effects on stimulus-evoked responses in visual cortex, we performed a probabilistic cueing experiment while recording electroencephalographic (EEG) data. Participants (n=48) learned associations between visual cues and subsequently presented gratings. A given cue predicted the appearance of a certain grating orientation with 10%, 25%, 50%, 75%, or 90% validity. We did not observe any stimulus expectancy effects on grating-evoked event-related potentials. Multivariate classifiers trained to discriminate between grating orientations performed better when classifying 10% compared to 90% probability gratings. However, classification performance did not substantively differ across any other stimulus expectancy conditions. Our findings provide very limited evidence for modulations of prediction error signalling by probabilistic expectations as specified in contemporary predictive coding models.

## 1. Introduction

Humans and other animals can readily learn associations between successive, co-occurring sensory events. They can exploit these statistical regularities to predict what is likely to occur in the immediate future. These predictions are thought to help us detect (and learn from) unusual or novel occurrences (Ulanovsky et al., 2003; Press et al., 2020) and rapidly respond to predictable stimuli (Gold & Stocker, 2017). This capacity fits into a broader framework of predictive processes that are enabled by a variety of mechanisms within the brain (reviewed in Teufel & Fletcher, 2020).

Several influential models based on hierarchical predictive coding (Friston, 2005, 2010; Bastos et al., 2012; Clark, 2013; Summerfield & de Lange, 2014; Keller & Mrsic-Flogel, 2018) describe effects of probabilistic expectations on stimulus-selective neurons in the visual system. According to these models, sensory systems are continuously attempting to infer the underlying causes of afferent sensory signals, whereby neurally instantiated models are continuously updated to best match patterns of sensory input. Discrepancies between these internal models and sensory input are thought to be indexed by activity of prediction error neurons (specifically, cortical pyramidal neurons) which are prevalent throughout the visual system (e.g., Friston, 2010; Bastos et al., 2012).

These predictive coding accounts posit that our internal models can be proactively adjusted in anticipation of expected sensory input, which allows a reduction of prediction error signalling when a stimulus matches our expectations, as compared to when that stimulus is unexpected (e.g., Egner et al., 2010; Summerfield & de Lange, 2014). This hypothesised effect has been termed expectation suppression (Summerfield et al., 2008; Todorovic & de Lange, 2012), and is related to the more general hypothesis that the responses of stimulus-selective neurons (and measured neural responses) will inversely scale with the subjective appearance probability of a presented stimulus (Walsh et al., 2020). Expectation suppression effects have been proposed to occur via *dampening*, whereby neurons that respond most strongly to an unpredicted stimulus are selectively suppressed when that stimulus is expected (Richter et al., 2022), congruent with reduced prediction error signalling. They have alternatively been proposed to occur via *sharpening*, whereby neurons that are not selective for a stimulus are suppressed within a given neural population, but there is an overall reduction in measured response magnitude (Kok et al., 2012; Yon et al., 2018). Notably, several other predictive processing-based accounts (which do not appeal to subtractive prediction error minimisation) also specify neural response increases for highly unexpected stimuli that are broadly in-line with this general hypothesis (e.g., Press et al., 2020; Alink & Blank, 2021; Marvan & Phillips, 2024).

The hypothesis described in Walsh et al. (2020) has been extensively tested using probabilistic cueing designs, whereby participants form expectations to see particular stimuli in specific contexts. In these designs, a visual stimulus is repeatedly presented following the same cue stimulus, so that the cue signals a high probability of that stimulus appearing. Stimuli that conform to these learned expectations are termed expected stimuli, and those that violate these expectations are termed surprising stimuli (following terminology in Egner et al., 2010; Rahnev et al., 2011; Amado et al., 2016). A subset of experiments also presented conditions whereby two or more different stimuli are each expected to appear with equal probability, termed neutral stimuli (for further definition see Arnal & Giraud, 2012; Feuerriegel et al., 2021). These neutral conditions have been included to isolate and quantify effects of fulfilled expectations (i.e., expectation suppression, using expected-neutral condition contrasts) and to separately quantify effects of expectation violations (using surprising-neutral contrasts). Please note that the terms ‘expected’, ‘surprising’ and ‘neutral’ are defined here in ways that are consistent with how these terms have been used in closely related work (e.g., Egner et al., 2010; Rahnev et al., 2011; Amado et al., 2016). These definitions may differ from how similar terms are used in other areas of the predictive processing literature. For example, in relation to broad expectations such as ‘some unspecified stimulus will appear’ all stimuli in the above examples could be reclassed as expected stimuli (Kwisthout et al., 2017). Here, we define a stimulus as expected if it is subjectively more likely to appear than the other stimulus identities that could possibly be presented.

Across probabilistic cueing studies that involved electrophysiological measures of neural activity, for example firing rates, local field potential amplitudes, or electroencephalographic (EEG) or magnetoencephalographic (MEG) signals, clear evidence for expectation suppression in the visual system has not been observed (Kaliukhovich & Vogels, 2011; Vinken et al., 2018; Rungratsameetaweemana et al., 2018; Solomon et al., 2021; reviewed in Feuerriegel et al., 2021; den Ouden et al., 2023). Although Kok et al. (2017) reported a pre-activation effect whereby expected stimuli could be decoded prior to stimulus onset based on patterns of MEG signals, they did not observe expectancy-related differences in the magnitude of grating stimulus-evoked event-related fields. Song et al. (2024) manipulated participants’ expectations about whether a stimulus would be repeated within a trial, or whether a different stimulus would subsequently appear. Repetition probability varied across blocks of trials. Although they observed repetition by expectation interaction effects on ERPs spanning approximately 330-630ms from stimulus onset, they did not find main effects of expectation. In other studies, expected-surprising condition differences have generally exhibited frontal or parietal topographic distributions that indicate sources outside of the visual system (Summerfield et al., 2011; Hall et al., 2018; Meijs et al., 2018; Feuerriegel et al., 2018; Alilovic et al., 2019). Probabilistic cueing effects have been reported in macaques within statistical learning experiments that involved weeks of sequence learning (Meyer & Olson, 2011; Meyer et al., 2014; Ramachandran et al., 2017; Schwiedrzik & Freiwald, 2017; Kaposvari et al., 2018; Esmailpour et al., 2023). However, similar effects were not consistently observed when stimulus appearance probabilities were learned over shorter time periods in comparable designs, in macaques (Kaliukhovich & Vogels, 2011; Vinken et al., 2018; Solomon et al., 2021) and in humans (Manahova et al., 2018; Zhou et al., 2020). Although fMRI BOLD signal differences between expected and surprising stimuli have been reported (e.g., Summerfield et al., 2008; Egner et al., 2010; Grotheer & Kovacs, 2015), this appears to be specifically due to signal increases that occur when stimuli are surprising (indexed by surprising-neutral condition differences), rather than a genuine suppression that occurs when an observer’s expectations are fulfilled (indexed by expected-neutral condition differences; reviewed in Feuerriegel et al., 2021). Such surprise-specific effects complicate the interpretation of expected-surprising BOLD signal differences. Other factors, such as pupil dilation following surprising events (O’Reilly et al., 2013), can influence BOLD signals in ways that can be misinterpreted as modulations of prediction error signalling (discussed in Richter & de Lange, 2019; den Ouden et al., 2023). Due to these issues, and the fact that prediction error signalling has been theoretically linked to pyramidal cell firing rates and local field potentials (e.g., Friston, 2005; Bastos et al., 2012), electrophysiological evidence is particularly important for determining the presence or absence of expectation suppression as implemented via prediction error minimisation.

Notably, there are aspects of existing studies that may limit the inferences that can be drawn from these null results. In Rungratsameetaweemana et al. (2018) participants could not form predictions for specific patterns of retinal input (i.e., specific images), which may have impaired detection of expectation effects. Other studies recruited samples that may not have been sufficient to detect small magnitude effects (e.g., n=23 in Kok et al., 2017; n=22 in Solomon et al., 2021).

To address these limitations, we previously recruited a large (n=48) sample and performed a probabilistic cueing experiment in which different face identities were presented (den Ouden et al., 2023). We also avoided known confounds that can mimic expectation suppression effects (reviewed in Feuerriegel et al., 2021). We did not observe any evidence for stimulus expectancy effects on face-evoked event-related potentials (ERPs) despite our large sample and extensive training of the cue-face associations in the experiment. However, we note that our findings may be specific to expectations about face identities, and that the face images did not differ markedly with respect to lower-level stimulus features (such as local luminance or shape). This precludes strong inferences about whether expectation effects are detectable in other neural populations, such as orientation-selective neurons in area V1 that have also been extensively investigated in fMRI probabilistic cueing designs (e.g., Kok et al., 2012).

To provide a more comprehensive evidence base spanning different stimulus types and populations of stimulus-selective neurons, we recruited a large (n=48) sample of participants who completed a similar probabilistic cueing experiment to that of den Ouden et al. (2023). In this study, cue images signalled the appearance probability of horizontally or vertically oriented gratings. Stimuli could appear with probabilities of 10%, 25% (labelled surprising stimuli), 50% (labelled neutral stimuli), 75%, or 90% (labelled expected stimuli). As researchers do not have direct access to participants’ internally generated expectations, we tested whether the learning of probabilistic cue-stimulus associations is sufficient for modulating EEG measures of stimulus-evoked responses in the visual system. We compared ERPs evoked by expected, neutral and surprising gratings using mass-univariate and multivariate pattern analyses (MVPA). We included the neutral condition to isolate and quantify effects associated with expectation fulfillment, assessed using expected-neutral contrasts, and expectation violation, assessed using surprising-neutral contrasts. This was done to test for any effects that are specific to higher or lower values of stimulus appearance probability, such as the surprise-related BOLD signal effects described above. The hypothesis in Walsh et al. (2020) does not specify a linear linking function between subjective probability and neural response magnitude, and so our approach allowed us to also identify non-linear relationships (discussed in Feuerriegel et al., 2021). As done in den Ouden et al. (2023), we also tested for stimulus repetition effects to compare their time-courses to any observed expectation effects. This was done to establish whether expectation and repetition effects show comparable or distinct time-courses, which may imply shared or distinct underlying mechanisms (reviewed in Feuerriegel, 2024).

To complement our ERP measures, we additionally trained multivariate classifiers to discriminate between different grating orientations using data collected during separate, randomised grating presentation blocks. These blocks involved gratings of varying orientations that were presented in a random (i.e., unpredictable) order. This allowed us to assess whether expectations influence distributed patterns of stimulus-selective responses by comparing decoding performance across expected, neutral and surprising stimulus conditions in the probabilistic cueing experiment (following Kok et al., 2017; Tang et al, 2018). In this context, dampening accounts based on expectation suppression and prediction error minimisation (Richter et al., 2022) would predict lower decoding performance for expected as compared to neutral and surprising stimuli. The opposite would be predicted by an alternative class of sharpening accounts of stimulus expectation effects (Kok et al., 2012; Yon et al., 2018; Press et al., 2020).

## 2. Methods

### 2.1. Participants

Fifty participants were recruited for this study. Participants were fluent in English and had normal or corrected-to-normal vision. One participant was excluded due to excessive, large amplitude artefacts detected in their EEG data. One participant was excluded due to poor task performance (less than 50% accuracy in the experimental task). This left 48 participants for both behavioural and EEG data analyses (35 women, 10 men, 3 other/non-binary, 46 right-handed) aged between 18 and 36 years (*M* = 24.9, *SD* = 4.7). We aimed to recruit a comparable sample size to den Ouden et al. (2023, n=48 included for analyses). As this study (and comparable previous work) did not find statistically significant expectation effects, an *a priori* effect size target could not be determined for statistical power analyses. However, our large sample size ensured high measurement precision for frequentist and Bayesian statistical tests. In this experiment, we would expect to observe moderate to large expectation effects on ERP amplitudes if the activity of superficial pyramidal neurons within visual cortex predominantly signals prediction errors that scale with subjective appearance probability (Friston, 2005; Bastos et al., 2012). Participants were reimbursed a gift voucher worth 45 AUD for their time. This study was approved by the Human Research Ethics Committee of the University of Melbourne.

### 2.2. Stimuli

Stimuli were presented at a distance of 80cm on a 27” Benq RL2755 LCD monitor (60Hz refresh rate) with a grey background in a darkened room. Stimuli were presented using MATLAB and functions from PsychToolbox v3 (Brainard, 1997; Kleiner et al., 2007). Code used for stimulus presentation will be available at osf.io/ygm69 at the time of publication.

Stimuli used during the experiment are depicted in Figure 1. All stimuli were presented concurrently with a fixation stimulus at the centre of the screen. The fixation stimulus consisted of a black circle (radius 0.90° of visual angle), a grey cross of the same luminance as the screen background, and a dot in the centre (Thaler et al., 2013). Cues and grating images were used for the training and probabilistic cueing blocks. Cue stimuli were coloured rings (1.22° inner edge radius, 0.36° width) presented around the fixation stimulus, which could be either blue, orange, or pink. Cues could either consist of a single thick ring, or two concentric rings separated by a spacer ring of the same colour as the background. Grating stimuli were circles (0.93° inner edge radius, 3.44° outer edge radius) that contained alternating black and white stripes (2.8 cycles/degree, example depicted in Figure 1E). Gratings in the training and probabilistic cueing blocks could be vertically or horizontally oriented.

**Figure 1.**
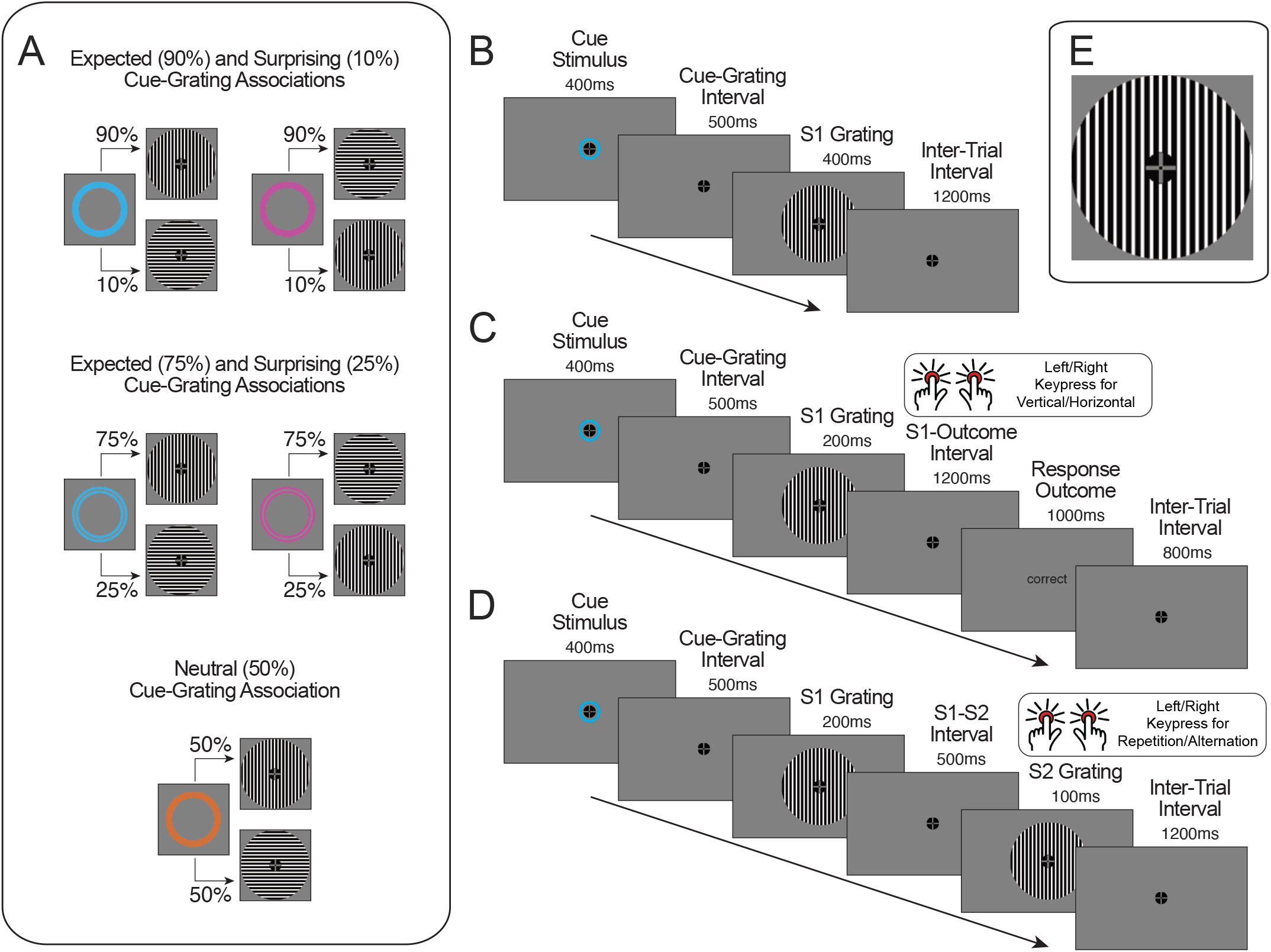
Cue-grating associations and trial diagrams. A) An example set of cue-grating associations for a participant. Each circle cue predicted the subsequent grating appearance of one of two gratings with 90% probability, 75% probability (expected stimuli), 50% probability (neutral stimuli), 25% probability or 10% probability (surprising stimuli). B) An example trial in the training session whereby participants were passively exposed to cue-grating pairs. C) An example trial from the training session grating orientation identification task. Participants were required to press one of two keys depending on the grating orientation presented in each trial. D) An example trial from the probabilistic cueing experiment. Participants were required to press one of two keys based on whether the second (S2) grating presented in the trial was the same or a different orientation to the first (S1) grating. Neither the cue identity or the S1 grating orientation were predictive of the S2 grating orientation being a repetition or alternation. E) An example vertical grating stimulus enlarged for easier visibility.

Gratings were also presented during separate randomised grating presentation blocks, which were included for training multivariate classifiers to discriminate between grating orientations. In these blocks, gratings could be one of eight orientations (22.5, 45, 67.5, 90, 112.5, 135, 157.5, or 180 degrees). Gratings presented during the randomised presentation blocks had spatial frequencies of 2.8 cycles/degree (non-targets) or 5.8 cycles/degree (targets).

### 2.3. Procedure

The experiment consisted of three phases that were completed sequentially: a training session, randomised grating presentation blocks, and a probabilistic cueing experiment. Prior to the experiment, the EEG recording equipment and cap were set up.

During the training session participants learned the probabilistic cue-grating associations (depicted in Figure 1A). The five cue-grating associations were introduced sequentially in separate sections. At the beginning of each section participants were shown a coloured circle cue image. They were then shown the associated gratings and explicitly told the appearance probability of each grating following that cue. For a given participant, one cue predicted a vertical grating with 90% probability (an expected stimulus), and a horizontal grating with 10% probability (a surprising stimulus). The cue of the same colour and different number of rings (one or two) predicted a vertical grating with 75% probability, and a horizontal grating with 25% probability. A different coloured pair of cues (including variants with one and two rings) predicted horizontal gratings as more likely with these same probabilities. This meant that the same-coloured cues predicted the same grating orientation as more probable, and the exact appearance probability depended on the number of rings visible in the cue. The remaining colour was assigned to the neutral condition, predicting a vertical grating with 50% probability and a horizontal grating with 50% probability. Only one variant (i.e., one or two rings) was used for a given participant as there was only one neutral condition. The neutral condition was included as a comparative ‘baseline’ condition to isolate and quantify effects of fulfilled expectations (i.e., expectation suppression) and surprise in our analyses. This allowed us to test for effects that are specific to differences across certain appearance probability values and identify any non-linear associations between neural response magnitude and subjective appearance probability (Walsh et al., 2020). The cue colours/numbers of rings and probabilistic associations with horizontal and vertical gratings were counterbalanced across participants.

Participants then passively viewed a block of trials during which a cue was presented for 400ms. After a 500ms interval a grating was presented for 400ms. Following the offset of this grating a fixation cross was presented for 1,200ms before the start of the following trial (Figure 1B). For each section (each cue), there were 20 of these example trials. Gratings followed the cues according to the instructed appearance probabilities. For example, for the cue predicting the vertical grating with 90% probability and horizontal grating with 10% probability, the vertical grating was presented eighteen times during the block and the horizontal grating was presented twice.

Within the training session, three sets of cues were presented to participants (order of sets counterbalanced across participants). These sets included the 90% vertical and 90% horizontal cues (set 1), the 75% vertical and 75% horizontal cues (set 2), and the 50% vertical/horizontal cue (set 3). The explicit cue-stimulus probability instructions and passive viewing were completed sequentially for each pair of cues within a set. For the set that only included the 50% neutral cue, the instructions and passive viewing trials were only presented for that cue.

After being exposed to the cue-grating associations within a set, participants completed an orientation identification task using only cues from that set. This task occurred after each set of cues were taught (i.e., 3 times in total). In each trial of this training session task, one of the cues were presented for 400ms, followed by a 500ms cue-stimulus interval and a grating presented for 200ms (Figure 1C). Participants were required to press keys ‘2’ or ‘5’ using their left or right index fingers on a TESORO Tizona numpad (1,000Hz polling rate) depending on which grating orientation (horizontal/vertical) was shown. Key assignments were counterbalanced across participants. Following their response in each trial, participants were provided feedback (‘correct’ or ‘error’) if their response was made within 1,200ms following grating onset. If their response was made before or after this time window, ‘too early’ or ‘too slow’ feedback was presented. Following the feedback, a fixation stimulus was presented for 800ms before the onset of the cue in the subsequent trial. Each grating orientation identification task block included 60 trials. This task served to strengthen the learned cue-grating associations and allowed us to determine whether participants had learnt the associations through analysing their task performance.

After each orientation identification task, participants were given a paper quiz with images of each cue and the different grating orientations with percentage probability values displayed above them. Participants drew lines to match the cue images they had just learned with the associated grating orientation appearance probabilities. The experimenter checked responses to ensure that cue-grating associations had been understood correctly. Any erroneous answers were corrected and explained to the participants. The training session lasted approximately twenty minutes.

After the training session participants completed the randomised presentation blocks, where they were presented with gratings of various orientations (22.5, 45, 67.5, 90, 112.5, 135, 157.5 and 180 degrees) and two different spatial frequencies (2.8 or 5.8 cycles/degree). In each trial a grating appeared for 200ms, followed by an inter-trial interval between 1,000-1,500ms (boxcar distribution, randomised across trials). Participants were required to identify target gratings (with a spatial frequency of 5.8 cycles/degree, presented in 8% of trials) by pressing ‘3’ on the numpad within 1,000ms after grating onset. They were instructed not to respond to non-target gratings (with a spatial frequency of 2.8 cycles/degree). Participants completed eight blocks in total, each including 100 trials. Following each block, participants were informed of their accuracy (proportion correctly identified targets) and average response time (RT) for correctly identified targets within that block. After completing this task, participants again completed the quiz that was completed during the training session to ensure that they could recall the associations prior to the probabilistic cueing experiment.

Participants then began the probabilistic cueing experiment. In each trial (depicted in Figure 1D) a cue was initially presented for 400ms. After a 500ms cue-stimulus interval, a first grating (termed the S1 grating) was presented for 200ms. After a 500ms interstimulus interval, a second grating (S2 grating) was presented for 100ms, followed by a 1,200ms inter-trial interval. Throughout all trials, the fixation stimulus was displayed at the center of the screen. Each cue stimulus was presented an equal number of times during the experiment. Probabilistic associations between cues and S1 gratings were the same as in the training session. Following each S1 grating, the S2 grating could either be a horizontal or vertical grating, each with 50% probability. Importantly, neither the cue or the S1 grating orientation were predictive of the S2 grating orientation, and participants were informed that no statistical association existed.

Participants completed a grating matching task during the probabilistic cueing experiment. Once the S2 grating appeared, participants were given 1,200ms to decide whether the S1 and S2 gratings were the same (repetitions) or different orientations (alternations). They did this by pressing keys ‘1’ and ‘3’ on the numpad with their left and right index fingers. Response key assignments were counterbalanced across participants. If participants did not make a response within 1,200ms after S2 grating onset, they received ‘too slow’ feedback for 1,000ms. If they made a response prior to S2 grating onset, ‘too fast’ feedback was presented. This task ensured that S1 gratings were task-relevant, but that no immediate decision or keypress response was required until S2 onset. This allowed us to avoid stimulus expectancy-related ERP effects associated with decision-making and motor response preparation (e.g., Gaillard, 1977; Steinemann et al., 2018; as done by Kok et al., 2017; den Ouden et al., 2023).

There were 100 trials per block, each consisting of 36 expected (90% probability) gratings, 30 expected (75% probability) gratings, 20 neutral (50% probability) gratings, 10 surprising (25% probability) gratings, and 4 surprising (10% probability) gratings. Participants completed six blocks (600 trials). After each block, participants were provided feedback on their accuracy and mean response times (RTs) for correct responses during that block. Participants took self-paced breaks between each block (minimum 10 seconds). The probabilistic cueing experiment (excluding breaks) lasted approximately 34 minutes.

Prior to the experiment, participants completed practice trials (minimum of 10) until they confirmed with the experimenter that they understood the task. Practice trials were identical to those in the probabilistic cueing experiment, however participants were provided with feedback (‘correct’, ‘error’, ‘too fast’, ‘too slow’) for 1,000ms after the response window following S2 grating onset in each trial. A fixation cross was displayed for 1,200ms between the offset of task feedback and the presentation of the cue in the subsequent trial.

### 2.4 Task Performance Analyses

To assess whether the probabilistic cue-grating associations were learned during the training session, we calculated participants’ accuracy (proportion correct) and mean RTs for trials with correct responses during the training session grating orientation identification task. Analyses were performed in JASP v0.16.4 (JASP Core Team). We compared performance between expected (90%) and surprising (10%) gratings, as well as expected (75%) and surprising (25%) gratings. These comparisons were selected because they involved trials that were intermixed within the same training session blocks. This allowed us to avoid the variability in within-subject difference measures that is associated with improvements in task performance that can occur across successive blocks of trials. As accuracy was near ceiling and not normally distributed, Wilcoxon signed-rank tests were used to test for differences in accuracy between conditions. Paired-samples t tests were used to test for differences in mean RTs. We used Bonferroni corrections to adjust alpha levels for multiple comparisons. We also used Bayesian versions of each test to derive Bayes factors in favour of the alternative hypothesis (Cauchy prior distribution, width 0.707, with 1,000 samples drawn for signed-rank tests).

Performance on the S1-S2 grating matching task during the probabilistic cueing experiment was measured by deriving accuracy and mean RTs for trials with correct responses. We compared accuracy across the five S1 appearance probability conditions using a Friedmans one-way ANOVA. We compared mean RTs across S1 probability conditions using a one-way repeated measures ANOVA (Greenhouse-Geisser correction). RTs were also compared across repeated and alternating S2 gratings (to assess stimulus repetition effects) using a paired-samples t test. Bayesian versions of the repeated measures ANOVA and paired-samples t test were also performed using JASP.

### 2.5. EEG Data Acquisition and Processing

We recorded EEG using a 64-channel Biosemi Active II system (Biosemi, The Netherlands) with a sampling rate of 512Hz using common mode sense and driven right leg electrodes (http://www.biosemi.com/faq/cms&drl.htm). We added six additional electrodes: two electrodes placed 1cm from the outer canthi of each eye, and four electrodes placed above and below the center of each eye.

We processed EEG data using EEGLab v2022.0 (Delorme & Makeig, 2004) in MATLAB (Mathworks) following the procedure in den Ouden et al. (2023). The EEG dataset as well as data processing and analysis code will be available at osf.io/ygm69 at the time of publication. First, we identified excessively noisy channels by visual inspection (mean excluded channels = 1.3, range 0-5) and excluded these from average reference calculations and Independent Components Analysis (ICA). Sections with large amplitude artefacts were also manually identified and removed. We then low-pass filtered the data at 40 Hz (EEGLab Basic Finite Impulse Response Filter New, default settings), referenced the data to the average of all channels and removed one extra channel (AFz) to compensate for the rank deficiency caused by average referencing. We duplicated the dataset and additionally applied a 0.1 Hz high-pass filter (EEGLab Basic FIR Filter New, default settings) to improve stationarity for the ICA. The ICA was performed on the high-pass filtered dataset using the RunICA extended algorithm (Jung et al., 2000). We then copied the independent component information to the non high-pass filtered dataset. Independent components associated with blinks and saccades were identified and removed according to guidelines in Chaumon et al. (2015). After ICA, we interpolated previously removed noisy channels and AFz using the cleaned dataset (spherical spline interpolation). EEG data were then high-pass filtered at 0.1 Hz.

The resulting data were segmented from −100 to 600ms relative to grating onset in the randomised presentation blocks, cue onset, S1 grating onset, and S2 grating onset. Epochs were baseline-corrected using the pre-stimulus interval. Epochs containing amplitudes exceeding ±100 µV from baseline at any of the 64 scalp channels, as well as epochs from trials with ‘too fast’ or ‘too slow’ responses to S2 gratings, were rejected (randomised presentation blocks non-target gratings: mean epochs retained = 629 out of 736, range 336-735, cues: mean epochs retained = 504 out of 600, range 297-576, S1 gratings: mean = 546, range 318-600, S2 gratings: mean = 548, range 326-600). To assess differences across cue types (which may also influence pre-S1 grating ERP baselines) in a supplementary analysis, we also segmented data from −100 to 900ms relative to cue onset. This included the 100ms window that was used for S1 grating epoch baseline subtraction.

### 2.6. Mass-Univariate Analyses of ERPs

As grating stimulus-evoked ERP components are prominent over occipital channels corresponding to populations of orientation-selective neurons (Lennie & Movshon, 2005; Roe et al., 2012), we defined an occipital region of interest (ROI) covering electrodes Oz, O1, O2, POz, and Iz. Trial-averaged ERPs in each condition were averaged across channels within this ROI prior to analyses. To provide more extensive coverage of the visual system we repeated these analyses using a parieto-occipital ROI spanning electrodes PO7/8, P7/8 and P9/10 (results in the Supplementary Material).

To test for expectation effects across the time-course of the S1 grating-evoked response, we used mass-univariate analyses of ERPs as in den Ouden et al. (2023). We performed paired-samples t tests at each time point relative to stimulus onset. Cluster-based permutation tests (based on the cluster mass statistic) were used to account for multiple comparisons (Maris & Oostenveld, 2007, including 358 comparisons per analysis, 10,000 permutation samples, cluster forming alpha = .01, family-wise alpha = .05) using functions from the Decision Decoding Toolbox v1.0.5 (Bode et al., 2019). For each permutation sample, condition labels were swapped for a random subset of participants. Paired-samples t tests were performed using this permuted-labels sample at each time point. All t values corresponding to p-values of < .01 were formed into clusters with any neighbouring such t values. Adjacent time points were considered temporal neighbours. Here, the sum of t values within each cluster is termed the ‘mass’ of that cluster. The largest cluster masses from each of the 10,000 permutation samples were used to estimate the null distribution. The masses of each cluster identified in the original dataset (with preserved condition labels) were then compared to this null distribution. The percentile ranking of each cluster mass value relative to the null distribution was used to derive the p-value for each cluster. This method provides control over the weak family-wise error rate while maintaining high sensitivity to detect effects in temporally-autocorrelated EEG data (Maris & Oostenveld, 2007; Groppe et al., 2011).

We performed separate mass-univariate analyses to test for each hypothesised effect. To determine whether combined effects of expectation suppression and surprise could be observed in our data, we compared S1-evoked ERPs from expected (90%) and surprising (10%) gratings, as well as expected (75%) and surprising (25%) gratings. To test for expectation suppression effects, we compared S1-evoked ERPs across expected (90%) and neutral (50%) gratings, as well as expected (75%) and neutral (50%) gratings. To test for surprise effects, we compared S1-evoked ERPs across surprising (10%) and neutral (50%) gratings, as well as surprising (25%) and neutral (50%) gratings.

To derive estimates of evidence for the null and alternative hypotheses we calculated Bayes factors at each time point using Bayesian paired-samples t tests as implemented in the BayesFactor toolbox v2.3.0 (Krekelberg, 2022, Cauchy prior distribution, width 0.707). Here, we caution that there are important issues relating to the use of Bayes factors for evaluating point null hypotheses, particularly when the alternative hypothesis spans a broad distribution of effect sizes (e.g., when using ‘pragmatic’ Cauchy prior distributions). In these cases, Bayes factors in favour of the null may simply indicate that the true effect size is too small to be detected and not necessarily zero, which is a limitation shared with frequentist approaches (Tendeiro & Kiers, 2019; Ly & Wagenmakers, 2022; Tendeiro et al., 2024). In the absence of specific alternative hypothesis effect size distributions that can be derived from existing theoretical models, we have chosen a Cauchy distribution as it is widely used in psychology and neuroscience (where BF_10_ > 1 corresponds to p < .05 in equivalent frequentist paired-samples t-tests). However, we do not interpret Bayes factors of BF10 < 1 as indicating explicit support for a point null hypothesis in this paper. Instead, we interpret this as indicating a lack of support for the alternative hypothesis.

We additionally tested for within- and across-trial image repetition effects using mass-univariate analyses as described above (following den Ouden et al., 2023). To measure within-trial repetition effects we compared ERPs evoked by S2 gratings when they were preceded by the same S1 grating orientation as opposed to a different orientation. To test for across-trial repetition effects we compared ERPs evoked by the coloured cues in cases where the cue in the previous trial was the same as compared to a different cue identity. We also compared ERPs evoked by S1 gratings, depending on whether the S2 grating in the previous trial was of the same or different orientation.

We also compared ERPs evoked by different cue types (90%, 75%, and 50% probability cues) using mass-univariate analyses applied to the −100 to 900ms epochs (results presented in the Supplementary Material).

### 2.7. Multivariate Pattern Classification Analyses

#### 2.7.1. Testing for ERP pattern differences across probabilistic cueing conditions

To complement the ROI-based ERP analyses described above, we additionally used MVPA to detect within-subject pattern differences that might qualitatively vary across individuals or occur outside of the predefined ROIs. We used support vector machine (SVM) classification as implemented in DDTBOX v1.0.5 (Bode et a., 2019) interfacing LIBSVM (Chang & Lin, 2011). We trained classifiers to discriminate between expected (90%) and surprising (10%) gratings, and also expected (75%) and surprising (25%) gratings, to determine if any combined effects of expectation and surprise were reflected in ERP pattern differences. These were preceded by the same cues in each trial, meaning that any late cue image-specific ERPs should not consistently produce above-chance classification performance (e.g., via effects on pre S1 grating ERP baselines). To be clear, in these analyses, we did not classify grating orientation, but rather the expected or surprising status of the gratings. We also used MVPA to test for ERP pattern differences across repeated and alternating S2 gratings (following den Ouden et al., 2023).

S1 and S2 grating-locked epochs were divided into non-overlapping analysis time windows of 10ms in duration. Within each time window, EEG amplitudes at each of the 64 scalp channels were averaged separately to create a spatial vector of brain activity (64 features) corresponding to each trial. In cases where epoch numbers were imbalanced across conditions, a random subset of epochs was drawn from the condition with more epochs to match the condition with fewer epochs. Data from each condition were then split into five subsets (i.e., five folds). SVM classifiers (cost parameter C = 1) were trained on the first four subsets from each condition and were subsequently tested using the remaining subset to derive a classification accuracy measure. This was repeated until each subset had been used once for testing (i.e., a five-fold cross-validation procedure). This cross-validated analysis was repeated another four times with different epochs randomly allocated to each subset to minimise effects of drawing biases. The average classification accuracy across all cross-validation steps and analysis repetitions was taken as the estimate of classification performance for each participant. This procedure was repeated with randomly permuted condition labels assigned to each trial to derive an empirical chance distribution.

Classification performance was compared between the original data and permuted-labels data at the group level using paired-samples t tests. As we aimed to detect above-chance classification performance, we used one-tailed tests. We used cluster-based permutation tests (settings as described above) to correct for multiple comparisons across analysis time windows.

#### 2.7.2. Decoding of grating orientations

In addition to testing for differences in distributed patterns of ERPs across probabilistic cueing conditions, we also trained classifiers to discriminate between grating orientations using data from the randomised presentation blocks. We then assessed whether classification accuracy differed across stimulus expectancy (for S1 gratings) and S1/S2 repetition or alternation conditions (for S2 gratings).

We first trained sets of classifiers using the epochs of EEG data corresponding to pairs of grating orientations in the randomised presentation blocks. We trained classifiers to discriminate between vertical grating stimuli and all other orientations (7 pairs of gratings orientations), as well as between the horizontal grating stimuli and all other orientations (7 pairs of grating orientations, for a total of 14 pairs). To avoid imbalances in numbers of exemplars across classes (which can lead biases in trained classifiers, Thölke et al., 2023), for each pair of grating orientations a randomly-selected set of trials was drawn from the condition with the higher number of retained epochs. SVM classifiers were then trained using the same parameters as described above, for each individual time point (sample) of data relative to stimulus onset (total of 358 time points). This resulted in a total of 14 trained classification models per time point per participant. Multiple classifiers (trained using different sets of grating orientations) were used to help minimise the variance in classification accuracy that is associated with idiosyncrasies of any single trained classifier model. This is expected to increase measurement precision at the participant level when classification performance is averaged across grating orientation sets.

We then tested classification performance using epochs that were time-locked to each S1 grating in the probabilistic cueing experiment. This was done for trials within each of the five S1 appearance probability conditions separately. Please note that, in the probabilistic cueing blocks, only vertical or horizontal gratings were presented to participants. Here, accuracy was defined as the proportion of trials in which the classifiers (trained on data from the randomised presentation blocks) correctly determined that the presented grating was a vertical grating (in trials with vertical gratings) or a horizontal grating (in trials with horizontal gratings), as opposed to the other grating orientation that a given classifier was trained to discriminate between.

The proportions of correctly classified gratings were then averaged across the different trained classifiers to produce a single metric of classification accuracy (e.g., as done by Hogendoorn & Burkitt, 2018; Blom et al., 2020). This metric captures the average proportion of instances that classifiers correctly predicted the (vertical or horizontal) grating orientation that was presented to the participant, as compared to the other grating orientations that appeared in randomised presentation blocks. Please also note that, because the classifiers are trained on equal numbers of exemplars from data in the randomised presentation blocks, there is not a risk of model training bias due to only one orientation being presented during the probabilistic cueing blocks.

Before comparing accuracy measures across probabilistic cueing conditions, we first verified that the trained classifiers could correctly identify the S1 gratings at above-chance levels. To do this, we averaged classification accuracy across all S1 probability conditions within each participant. We also repeated the same analyses but with the condition labels being randomly permuted in the training datasets. We then compared classification performance in the original data against permuted-labels classification performance using one-tailed, single sample t tests and cluster-based permutation tests and Bayesian t tests as described above. Following the structure of our ERP analyses, we then compared classification accuracy across 90% and 10% conditions, 75% and 25% conditions, 90% and 50% conditions, 75% and 50% conditions, 25% and 50% conditions, and 10% and 50% conditions using cluster-based permutation tests and Bayesian paired-samples t tests.

We also performed the above analyses using S1 grating-locked epochs that were baseline-corrected using the 100ms period before the onset of the cue stimulus in each trial. This was done because, in conditions where a specific grating orientation was expected, EEG responses may have been generated during the pre S1 grating baseline period that resembled the visual evoked responses following presentation of the expected grating (known as pre-activation, Kok et al., 2017). If these patterns were indeed generated (and were subtracted from the data via the baseline correction procedure) then classification performance might be artefactually lower in expected stimulus conditions over post S1 grating time windows where the same orientation-selective EEG patterns are normally elicited by the grating stimuli. Using pre-cue baselines in supplementary analyses allowed us to avoid this potential confound. Results are presented in the Supplementary Material.

To also assess stimulus repetition effects on classification performance (to compare to any stimulus expectation effects), we also tested the classifiers using EEG data time-locked to the repeated and alternating S2 gratings using the analysis methods described above.

We also performed post-hoc, exploratory temporal generalisation analyses (King & Dehaene, 2014) to assess whether classifiers trained using data at a given time point would be able to successfully classify grating orientations at different time points relative to stimulus onset. These additional analyses are described in the Supplementary Material.

## 3. Results

### 3.1. Training Session Task Performance

Accuracy and mean RTs for each appearance probability condition in the grating orientation identification task are plotted in Figure 2A-B. Accuracy was on average higher, and RTs were faster, for expected and neutral compared to surprising gratings. Participants achieved higher accuracy for expected (90%) compared to surprising (10%) gratings, *Z* = 3.45, *p* < .001, BF_10_ = 135.78, and expected (75%) compared to surprising (25%) gratings, *Z* = 2.71, *p* = .007, BF_10_ = 12.14 (Figure 2A). RTs were faster for expected (90%) compared to surprising (10%) gratings, *t*(47) = −4.49, *p* < .001, BF_10_ = 475.48, and expected (75%) compared to surprising (25%) gratings, *t*(47) = −2.72, *p* = .009, BF_10_ = 4.16 (Figure 2B). Higher accuracy and faster RTs for expected gratings indicates that participants successfully learned the cue-grating associations during the training session.

**Figure 2.**
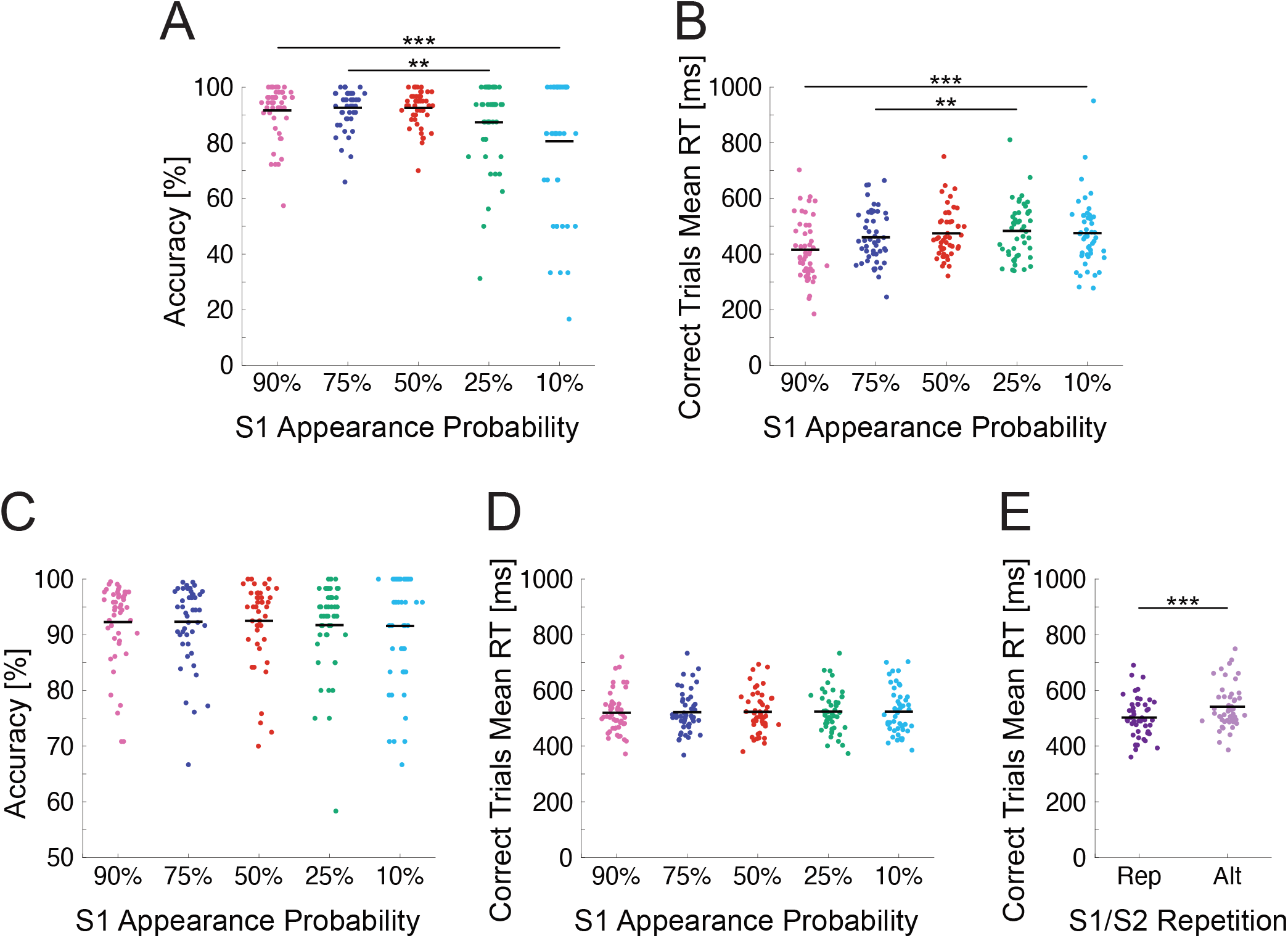
Task performance during the training session and grating matching task. A) Accuracy in the training session grating orientation identification task by cued stimulus appearance probability. Dots represent values for individual participants. Horizontal black lines denote group means. B) Mean RTs for correct trials. Accuracy was higher (and RTs faster) for expected (90%, 75% appearance probability) compared to surprising (10%, 25%) stimulus conditions. C) Accuracy in the grating matching task by S1 grating appearance probability condition. D) Mean RTs for correct trials by S1 grating appearance probability. E) Mean RTs for correct trials by S1-S2 repetition (Rep) or alternation (Alt) status. Accuracy and RTs did not substantially differ across S1 probability conditions but RTs were faster for repeated compared to alternating S2 gratings. Lines with asterisks denote statistically significant differences between compared conditions (** denotes *p* < .01, *** denotes *p* < .001).

### 3.2. Probabilistic Cueing Experiment Task Performance

Accuracy was generally high across all S1 grating expectancy conditions during the S1-S2 grating matching task (Figure 2C). A Friedmans one-way ANOVA did not reveal differences between S1 appearance probability conditions, χ^2^ (4) = 3.12, *p* = .539, Kendall’s W = .02. Differences in mean RTs were not observed across stimulus appearance probability conditions, *F*(2.52, 118.38) = 0.59, *p* = .592, BF_10_ = 0.03 (Figure 2D). However, RTs for repeated gratings were faster than those for alternating gratings, *t*(47) = −8.49, *p* < .001, BF_10_ = 2.05 * 10^8^ (Figure 2E, consistent with Rostalski et al., 2020; den Ouden et al., 2023).

### 3.3. Expectation Effects on S1 Grating-Evoked ERPs

We did not find any statistically significant differences in ERP amplitudes across any of the compared S1 expectancy conditions. Figure 3 displays the group average ERPs for each set of conditions, the difference waves, standardised Cohen’s d effect size estimates and Bayes factors. We did not observe differences between ERPs evoked by expected (90%) and surprising (10%) gratings (Figure 3A), expected (75%) and surprising (25%) gratings (Figure 3B), expected (90%) and neutral (50%) gratings (Figure 3C), expected (75%) and neutral (50%) gratings (Figure 3D), neutral (50%) and surprising (10%) gratings (Figure 3E) and neutral (50%) and surprising (25%) gratings (Figure 3F). Observed amplitude differences and standardized effect sizes were generally very small, and Bayes factors generally indicated a lack of support for the alternative hypothesis throughout the time-course of the S1 grating-evoked response. Results of analyses of ERPs at the parieto-occipital ROI (included in the Supplementary Material) were also consistent with those described here.

**Figure 3.**
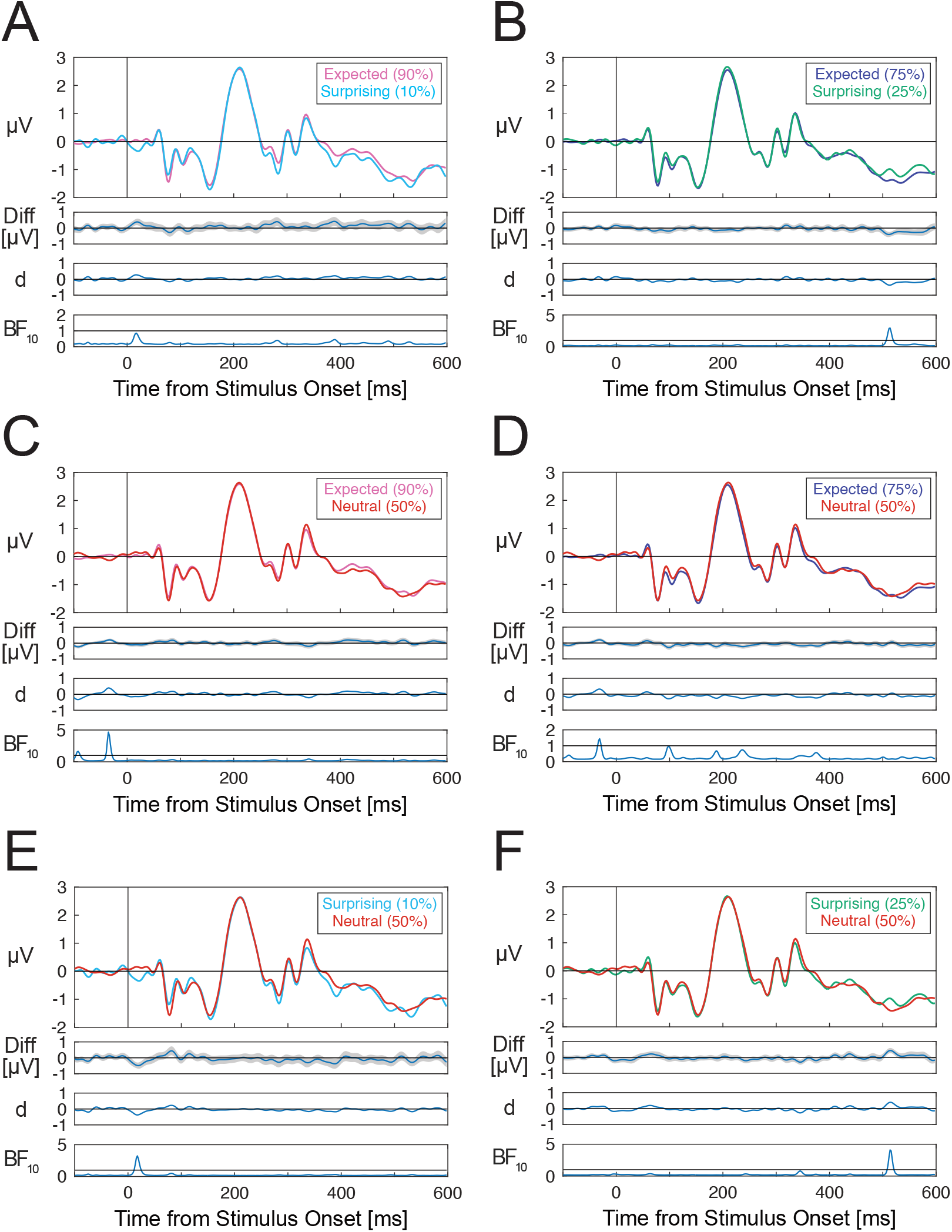
Group-averaged ERPs evoked by S1 gratings in different expectation conditions. A) Expected (90%) - surprising (10%) ERP differences. B) Expected (75%) - surprising (25%) differences. C) Expected (90%) - neutral (50%) differences. D) Expected (75%) - neutral (50%) differences. E) Neutral (50%) - surprising (10%) differences. F) Neutral (50%) - surprising (25%) differences. No statistically significant differences across S1 probability conditions were observed after correction for multiple comparisons. ERPs are averaged across channels within the occipital ROI including channels Oz/1/2, POz and Iz. ERPs for each set of compared conditions are displayed along with difference waves (with shading denoting standard errors), Cohen’s d estimates and Bayes factors. Horizontal lines in Bayes factor plots denote BF_10_ = 1, indicating a lack of preferential support for either the alternative or null hypothesis.

To verify whether cue-evoked ERPs differed across cue types (where such differences may influence pre-S1 grating ERP baselines), we also compared ERPs evoked by 90%, 75%, and 50% probability cues. We did not observe evidence for any ERP amplitude differences across cue types (results provided in the Supplementary Material).

### 3.4. Effects of Within-Trial Repetition on S2 Grating-Evoked ERPs

We observed several within-trial grating repetition effects between 76-600ms from S2 grating onset. Group-averaged ERPs for repeated and alternating S2 gratings are shown in Figure 4A. Scalp maps of effect topographies are plotted in Figure 4B. Four effects were found: between 76-113ms (cluster *p* = .013), between 150-176ms (cluster *p* = .044), between 357-492ms (cluster *p* < .001), and between 556-600ms from S2 grating onset (cluster *p* = .033). Effects over similar time windows were also observed at parieto-occipital electrodes (shown in the Supplementary Material).

**Figure 4.**
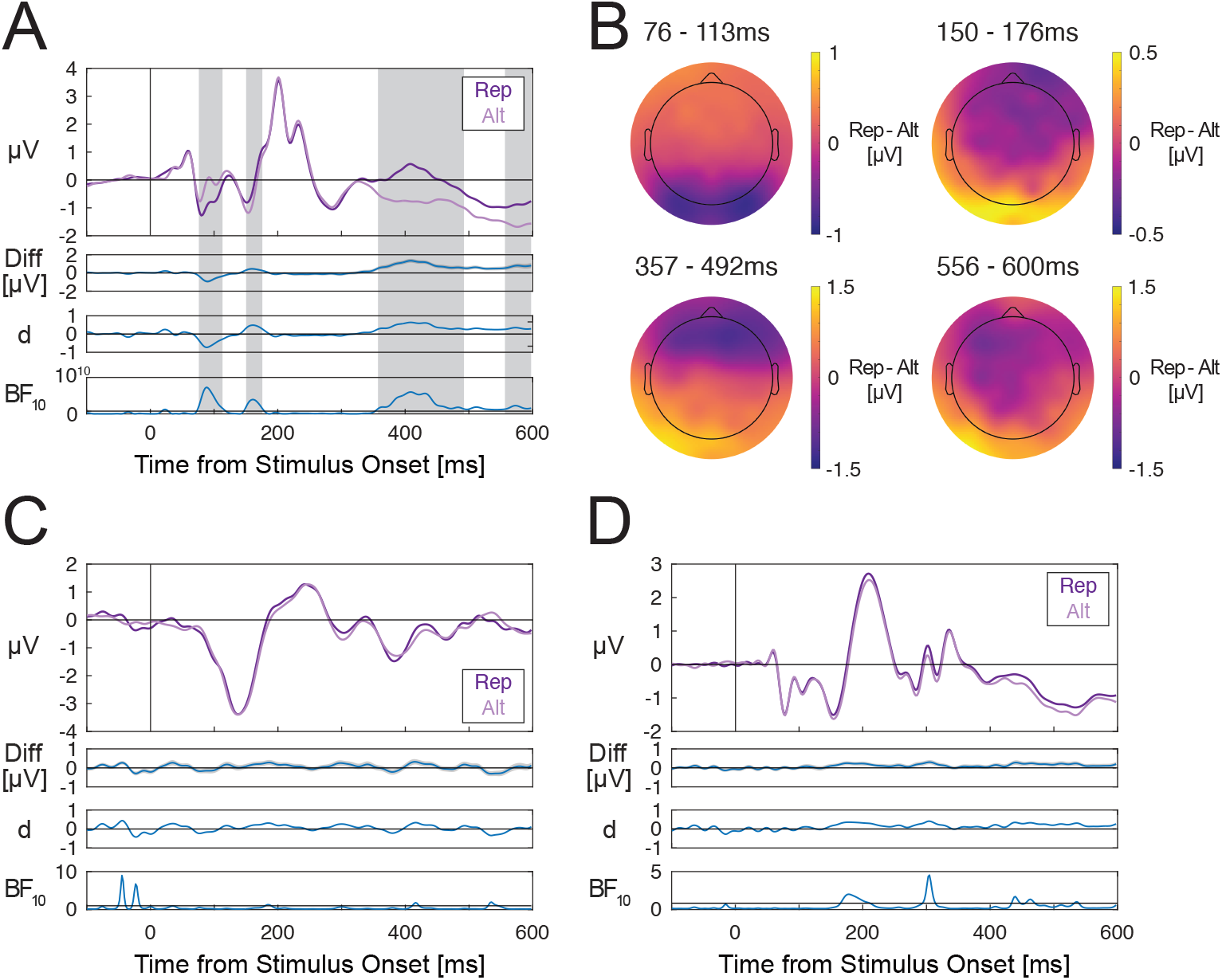
Within- and across-trial repetition comparisons for S2 gratings, cues, and S1 gratings. A) S2 grating-evoked ERPs depending on whether the S2 grating was the same orientation as the S1 grating (repetition/Rep) or a different orientation (alternation/Alt). B) Scalp maps of grating repetition effects for S2 gratings averaged over each statistically significant time window. C) Cue-evoked ERPs depending on whether the cue image was the same image as the cue in the previous trial (Rep) or a different cue (Alt). D) S1 grating-evoked ERPs depending on whether the S2 grating in the previous trial was the same (Rep) or a different orientation (Alt). There were clear within-trial (but not across-trial) repetition effects during several time windows spanning 79-600ms. ERPs are averaged across channels within the occipital ROI including channels Oz/1/2, POz and Iz. For each pair of compared conditions, ERPs are displayed along with difference waves (grey shading denoting standard errors), Cohen’s estimates and Bayes factors in favour of the alternative hypothesis. Note that for A) Bayes factors are plotted on logarithmic scales due to the wide ranges of values across the time-course of the stimulus-evoked response. Horizontal lines in Bayes factor plots denote BF10 = 1, indicating a lack of preferential support for either the alternative or null hypothesis. Grey shaded areas denote time windows of statistically significant differences after correcting for multiple comparisons.

### 3.5. Effects of Across-Trial Repetition on Cue- and S1 Grating-Evoked ERPs

We did not observe any ERP differences across cues that were the same as compared to different to the cue in the preceding trial (Figure 4C). We also did not find differences in ERPs evoked by S1 gratings that were the same compared to different to the S2 grating presented in the previous trial (Figure 4D). This was broadly consistent with results for parieto-occipital ERPs, except that two across-trial grating repetition effects were observed at those electrodes (reported in the Supplementary Material).

### 3.6. Multivariate Pattern Classification Results

We used MVPA to test for patterns of single-trial ERPs that discriminate between expected and surprising S1 gratings. S1 grating expectancy could not be classified at above-chance level at any time points, for expected (90%) and surprising (10%) gratings (Figure 5A), and for expected (75%) and surprising (25%) gratings (Figure 5B). We did not use MVPA to investigate other expectancy conditions, as the preceding cue images systematically differed across these conditions within each participant dataset. Late cue-specific ERP effects could potentially result in spurious above-chance classification by influencing S1 grating pre-stimulus baselines (see Feuerriegel & Bode, 2022).

**Figure 5.**
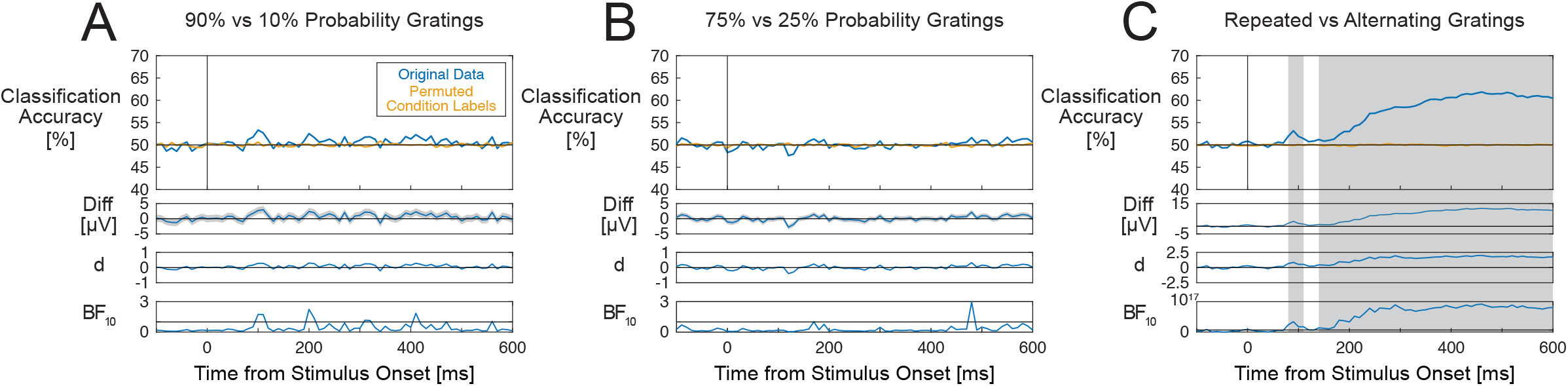
Multivariate pattern classification results. A) Classification accuracy for discriminating between expected (90% appearance probability) and surprising (10% appearance probability) S1 gratings. Blue lines display classification accuracy for the original data and orange lines show classification accuracy for the data with permuted condition labels (empirical chance distribution). B) Classification accuracy for discriminating between expected (75%) and surprising (25%) probability S1 gratings. C) Classification accuracy for discriminating between repeated and alternating S2 gratings. Classifiers could discriminate at above-chance levels between repeated and alternating S2 gratings, but not between different S1 grating appearance probability conditions. Grey shaded areas denote time windows where classification performance was statistically significantly above-chance after correcting for multiple comparisons. For each plot, classification accuracy is displayed along with difference waves (grey shading denoting standard errors), Cohen’s estimates and Bayes factors in favour of the alternative hypothesis. Note that for C) Bayes factors are plotted on logarithmic scales due to the wide ranges of values across the time-course of the stimulus-evoked response. Horizontal lines in Bayes factor plots denote BF10 = 1, indicating a lack of preferential support for either the alternative or null hypothesis. Grey shaded areas denote time windows of statistically significant differences after correcting for multiple comparisons.

By contrast, classifiers could differentiate between the repeated and alternating S2 gratings at above-chance levels between 80-110ms (cluster *p* = .010). and 140-600ms after S2 grating onset (cluster *p* < .001, Figure 5C).

### 3.7. Decoding Orientations of S1 and S2 Gratings

We also assessed whether stimulus expectations influence distributed, grating orientation-selective patterns of EEG signals, as indicated by changes in multivariate classification accuracy (Kok et al., 2017; Tang et al., 2018).

We first verified that classifiers trained using grating-evoked ERPs from the randomised presentation blocks could successfully discriminate between S1 grating orientations in the probabilistic cueing blocks. Above-chance decoding was observed across four time windows spanning approximately 50-300ms from S1 grating onset (Figure 6A, similar to Grootswagers et al., 2024).

**Figure 6.**
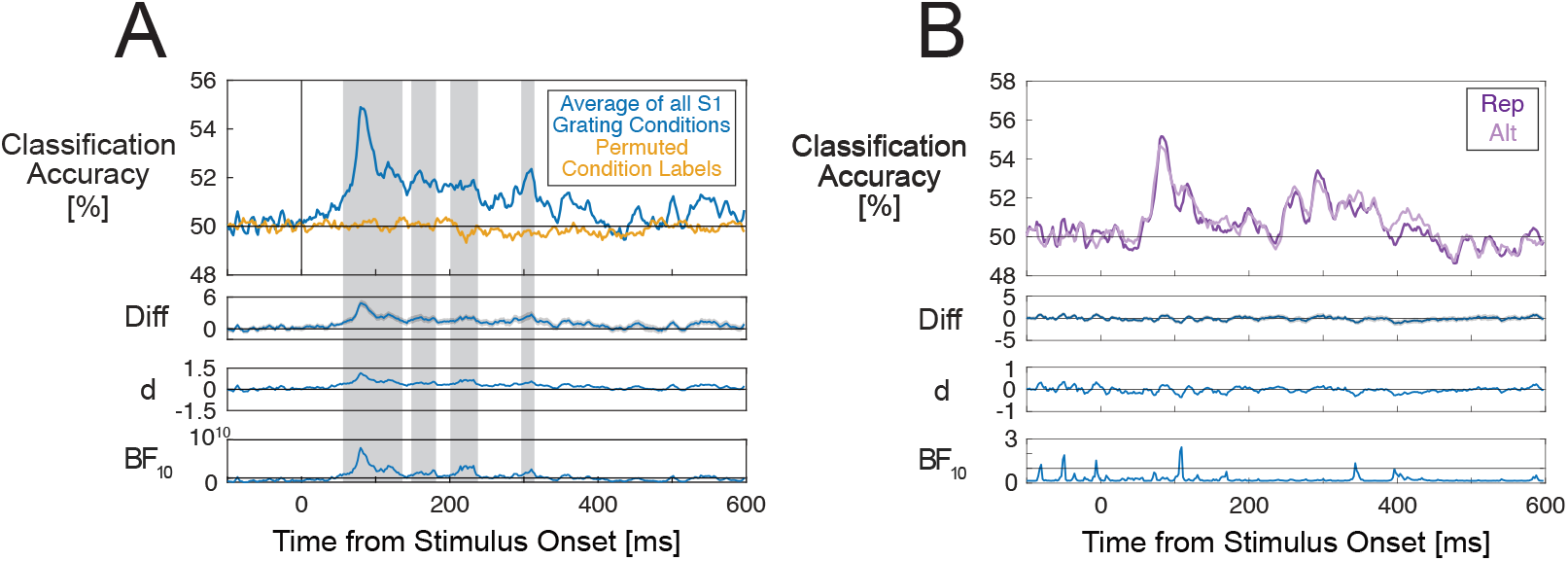
Classification of grating orientations for S1 and S2 stimuli. A) Classification accuracy for each time point relative to S1 grating onset, averaged across each S1 appearance probability condition. The blue line displays classification accuracy for the original data and the orange line shows classification accuracy for the data with permuted condition labels (empirical chance distribution). S1 gratings could be classified above chance at time windows spanning 50-300ms. Grey shaded areas denote time windows where classification performance was significantly above-chance after correcting for multiple comparisons. B) Classification performance for repeated and alternating S2 gratings. No statistically significant differences in classification performance were observed. Classification accuracy is displayed along with difference waves (grey shading denoting standard errors), Cohen’s estimates and Bayes factors in favour of the alternative hypothesis. For A) Bayes factors are plotted on a logarithmic scale due to the wide ranges of values across the time-course of the stimulus-evoked response. Horizontal lines in Bayes factor plots denote BF10 = 1, indicating a lack of preferential support for either the alternative or null hypothesis.

We then compared classification performance across different S1 grating appearance probability conditions. After correcting for multiple comparisons, we observed higher classification performance for surprising (10% appearance probability) as compared to expected (90% probability) conditions over a narrow time window spanning 281-302ms from S1 grating onset (cluster *p* = .013, Figure 7A). This appeared to be due to differences in accuracy between the 10% probability condition and other conditions. Other comparisons between 75% and 25%, 90% and 50%, 75% and 50%, 50% and 25%, and 50% and 10% probability conditions did not yield clear differences in classification performance that survived correction for multiple tests (Figure 7B-F). This is despite very brief time windows over which Bayes factors were higher than one, as can occur by chance when there are large numbers of tests. Over an earlier time window spanning approximately 150-2ooms there appeared to be higher accuracy for surprising (10%) compared to expected (90%) gratings, but this did not survive the multiple testing correction. When analyses were repeated using epochs that were baseline-corrected using the pre-cue period, no differences in classification performance were observed across S1 probability conditions (results presented in the Supplementary Material).

**Figure 7.**
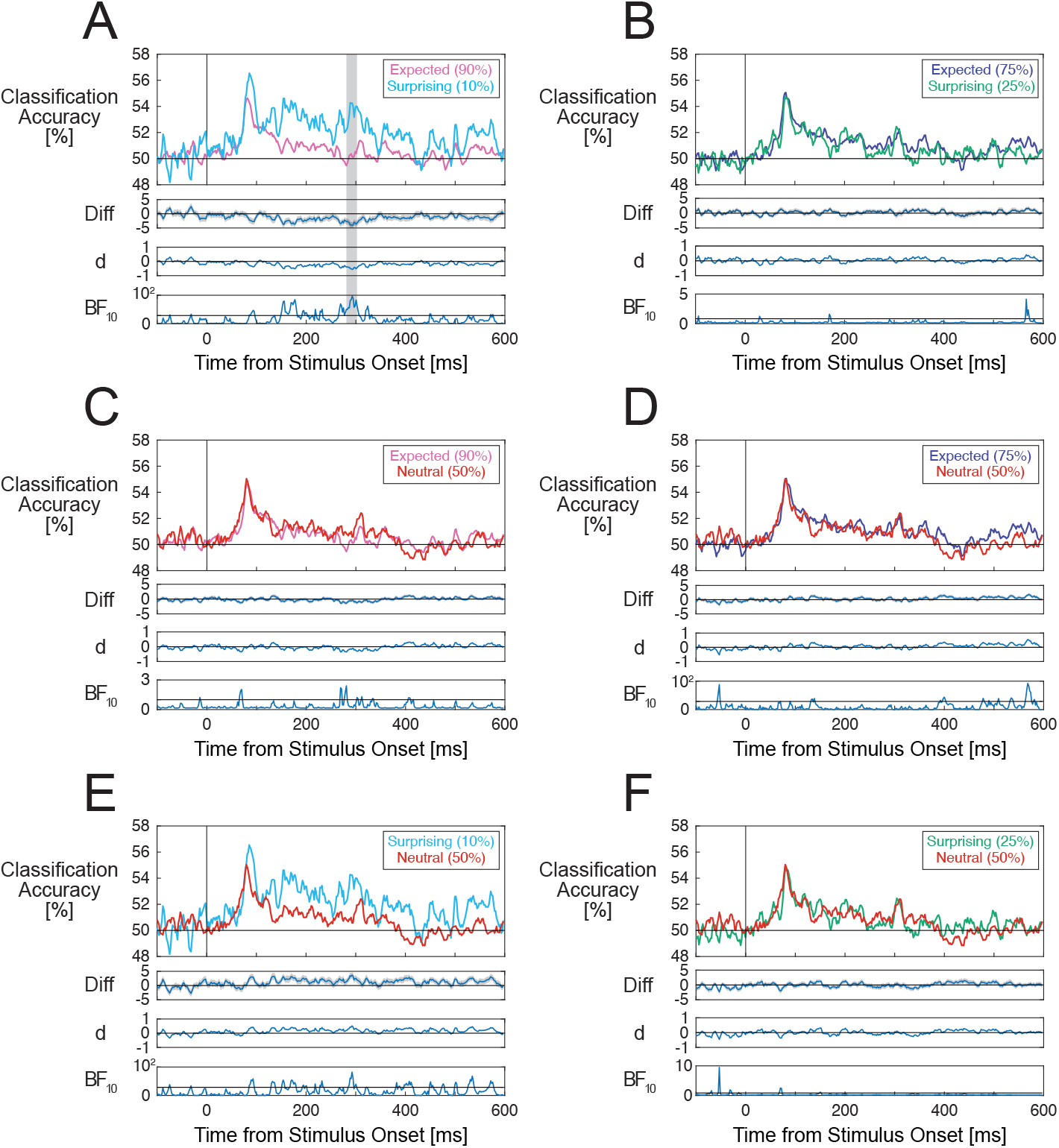
Classification performance results for classifiers trained to discriminate between S1 grating orientations. A) Classification performance for expected (90%) and surprising (10%) S1 grating conditions. B) Classification performance for expected (75%) and surprising (25%) conditions. C) Classification performance for expected (90%) and neutral (50%) conditions. D) Classification performance for expected (75%) and neutral (50%) conditions. E) Classification performance for surprising (10%) and neutral (50%) conditions. F) Classification performance for surprising (25%) and neutral (50%) conditions. Only differences between 90% and 10% probability conditions (over a time window spanning 281-302ms) survived correction for multiple comparisons. For each pair of conditions, classification accuracy is displayed along with difference waves (shading denoting standard errors), Cohen’s d effect size estimates, and Bayes factors in favour of the alternative hypothesis. Grey shaded areas denote time windows of statistically significant differences after correcting for multiple comparisons. Note that for A), D), and E) Bayes factors are plotted on logarithmic scales due to the wide ranges of values across the time-course of the stimulus-evoked response. Horizontal lines in Bayes factor plots denote BF10 = 1, indicating a lack of preferential support for either the alternative or null hypothesis.

As a comparison analysis, we also assessed whether classification accuracy differed across repeated and alternating S2 gratings. We did not observe any statistically significant differences (Figure 6B).

Post-hoc temporal generalisation analyses (presented in the Supplementary Material) did not show clear evidence of off-diagonal above chance classification performance beyond what would be expected based on the low-pass filter applied during EEG data processing. We also did not observe evidence for differences in classification performance across S1 probability conditions.

## 4. Discussion

We performed a probabilistic cueing experiment while recording EEG to provide a strong test for stimulus expectation effects on electrophysiological responses in the visual system. Using a similar experiment design to den Ouden et al. (2023), we trained participants (n=48) to associate specific cue stimuli with different probabilities (ranging from 10-90%) that a horizontal or vertical grating would subsequently appear. We verified that participants had learned these cue-stimulus associations during the training session, however we did not find evidence that the learning of these probabilistic cue-stimulus associations led to modulations of grating-evoked ERPs. Effect magnitude estimates were very small and Bayes factors generally did not favour the alternative hypothesis across the time-course of the grating-evoked response. By comparison, we observed clear effects of within-trial stimulus repetition. When we trained classifiers to discriminate between grating stimulus orientations using data from separate randomised presentation blocks, we also did not observe any consistent differences in grating orientation classification performance by stimulus expectancy. Our findings do not provide support for the idea that probabilistic cueing is sufficient to produce expectation suppression or other changes in the electrophysiological responses of stimulus-selective neurons within the visual system.

### 4.1. Effects of Expectations on Grating-Evoked ERPs

Consistent with our previous study in which face stimuli were presented (den Ouden et al. 2023), we did not find evidence that grating-evoked ERPs were modulated by stimulus expectations. Mass-univariate ERP amplitude analyses did not yield any differences that survived correction for multiple comparisons. Effect estimates were generally very small (i.e., less than ±0.5µV). Although there were very short windows over which Bayes factors favoured the alternative hypothesis, these are not necessarily indicative of genuine ERP effects (which typically span longer time periods), especially when considering the large numbers of tests that were performed. Generally, Bayes factors indicated a lack of support for the alternative hypothesis throughout the time-course of the grating-evoked response. Multivariate classifiers, which can exploit subtle differences in distributed patterns of ERPs across the scalp, also could not distinguish between expected and surprising stimulus conditions.

These findings are consistent with recent electrophysiological work that has reported a lack of expectation effects in similar probabilistic cueing designs, particularly when controlling for confounding factors that can mimic hypothesised expectancy effects in the visual system (Kok et al., 2017; Rungratsameetaweemana et al., 2018; Kaliukhovich & Vogels, 2011; Vinken et al., 2018; Solomon et al., 2021; reviewed in Feuerriegel et al., 2021; den Ouden et al., 2023). Although some studies that included expectancy manipulations within repetition probability, statistical learning and spatial cueing designs have reported effects of expectations on ERPs, these have been most prominent at central or frontal electrodes, rather than having clear sources within the visual system (Summerfield et al., 2011; Feuerriegel et al., 2018; Hall et al., 2018; Meijs et al., 2018; Alilovic et al., 2019). In contrast to our findings, Tang et al. (2018) reported expectation effects between 75-150ms post grating onset using cluster-based permutation tests. Although Walsh et al. (2024) did observe differences between expected and surprising grating stimuli with respect to steady state visual evoked potential (SSVEP) amplitudes over visual cortex, this emerged only after multiple testing sessions (and thousands of trials). Moreover, fulfilled expectations appeared to enhance rather than suppress visual evoked responses (i.e., the opposite of expectation suppression). Notably, these SSVEP effects were measured over a late time window spanning 680-975ms following the expected or surprising stimulus change, where participants had already reported the identity of the grating stimulus before the start of this measurement window in a substantial portion of trials.

When considered alongside the work reviewed above, the findings from our well-powered study provide further evidence that probabilistic cueing effects on electrophysiological measures are either remarkably small in magnitude so as to be undetectable, or absent during time windows over which perceptual inferences are typically formed. This evidence does not support the hypothesis specified in Walsh et al. (2020) that neural response magnitudes will inversely scale with the subjective appearance probability of a stimulus, which is implicit in other predictive coding accounts (e.g., Friston, 2005, 2010; Bastos et al., 2012; Clark, 2013; Summerfield & de Lange, 2014; Keller & Mrsic-Flogel, 2018).

Our findings also raise the question of whether it is necessary to appeal to notions of prediction error minimisation via expectation suppression to explain visual system function (discussed in Spratling, 2017; Mikulasch et al., 2023; Marvan & Phillips, 2024). As described in Teufel and Fletcher (2020), the visual system can exploit many different features of sensory environments, and some processes do not rely on so-called ‘top-down’ expectations or generative models instantiated within visual cortex. Learned cue-stimulus associations may influence neural activity and behaviour via other mechanisms that are distributed throughout the brain. Outside of the visual system, preparatory motor activity enables faster responses when a stimulus (and associated action) can be predicted in advance (de Lange et al., 2013; Gold & Stocker, 2017; Kelly et al., 2021; Walsh et al., 2024). Covariation of neural responses with reward prediction errors and information gain has also been reported in areas outside of the visual system, such as the insula and striatum (e.g., Loued-Khenissi & Preuschoff, 2020; Schultz, 2016).

In addition, there are well-documented effects of attention, which is often guided by expectations about which stimulus locations and/or features are likely to be informative or relevant to a task at hand (Corbetta et al., 1990; Reynolds & Heeger, 2009; Carrasco et al., 2011; Heeger, 2017; Gottlieb, 2023). In some cases, our attention can also be involuntarily directed toward surprising, salient, or novel stimuli (Press et al., 2020; Alink & Blank, 2021; Marvan & Phillips, 2024). Notably, these types of effects are predominantly theorised to be enabled via excitatory feedback connections to stimulus-selective visual neurons (e.g., Desimone & Duncan, 1995; Reynolds & Chelazzi, 2004; Moore, 2006; Herrmann et al., 2010) rather than inhibition (for comparisons of inhibition- and excitation-based predictive coding algorithms see Spratling, 2017; Marvan & Phillips, 2024). They are also distinct from the expectancy effects tested for in our experiment, whereby all S1 gratings were equally task-relevant. Although stimulus expectations and attention may be hypothesised to produce similar effects on neural response measures, they differ regarding their underlying mechanisms and the experimental contexts in which their effects will be observed (discussed in Rungratsameetaweemana & Serences, 2019; Alink & Blank, 2021).

The lack of expectation effects observed in electrophysiological measures contrasts with consistent observations of fMRI BOLD signal increases when stimuli are subjectively surprising, as compared to expected or neutral stimulus conditions (Summerfield & Koechlin, 2008; Amado et al., 2016; Richter & de Lange, 2019; reviewed in Feuerriegel et al., 2021). The interpretation of these BOLD signal effects as reductions of prediction error signalling (in the absence of comparable electrophysiological evidence) is complicated by other phenomena which may co-occur with surprise, which may influence BOLD signal magnitudes in visual cortex. These include pupil dilation following subjectively surprising events (O’Reilly et al., 2013) as well as time-on-task effects that covary with response times (Yarkoni et al., 2009; Mumford et al., 2024) and effects of attentional capture following unexpected stimuli (Press et al., 2020; Alink & Blank, 2021; for further discussion see Richter & de Lange, 2019; Feuerriegel et al., 2021; den Ouden et al., 2023). Further work is needed to characterise the contributions of these phenomena to expected-surprising stimulus BOLD signal differences (following Richter & de Lange, 2019).

Here, we also note that our findings do not challenge predictive coding model variants that do *not* specify expectation suppression effects (e.g., Rao & Ballard, 1999). However, many such models often provide indistinguishable predictions from so-called ‘classical’ accounts of visual system function (discussed in Aitchison & Lengyel, 2017; Cao, 2020; Williams, 2022).

### 4.2. Expectation Effects on Grating Orientation Decoding Performance

To additionally assess whether stimulus expectations alter distributed patterns of activity across stimulus-selective neurons, we trained multivariate classifiers to discriminate between gratings of different orientations using data from the randomised presentation blocks. These classifiers were then used to classify the orientations of S1 gratings that appeared with different cued probabilities. Although the classifiers could discriminate between S1 grating orientations at above-chance levels, we only observed higher classification performance for the surprising (10%) compared to the expected (90%) condition over a narrow time window (281-302ms post stimulus onset). We also observed higher classification accuracy during an earlier window spanning approximately 150-200ms, consistent with Tang et al. (2018) who observed larger orientation selective signals for low probability stimuli between 79-185ms post grating stimulus onset. However, this effect did not survive correction for multiple comparisons in our study, and differences were not observed for other sets of S1 probability conditions.

Probabilistic cueing effects on multivariate classification performance (or comparable metrics) have been inconsistent across studies. Kok et al. (2017) reported larger grating orientation-selective signals for expected as compared to surprising grating orientations between −40 and 530ms relative to grating onset (when training on signals during a time window spanning 120-160ms), whereas the opposite was reported between 79-185ms post grating onset in Tang et al. (2018). Lower classification accuracy was reported for improbable object images as compared to randomly ordered ones within rapid serial visual presentation streams in Moore et al. (2024), whereas no differences in classification performance were observed within streams of letter stimuli in Meijs et al. (2019). This evidence is from a mixture of MEG and EEG recordings, and from a variety of different multivariate analysis techniques and experiment designs that are all proposed to manipulate participants’ expectations. Replication studies will be necessary to gain a stronger understanding of when (and if) stimulus expectations alter multivariate classification performance.

More generally, the inferences drawn from observed changes in classification performance depend on several key assumptions. If assuming that grating-evoked ERPs in the randomised presentation blocks (used to train classifiers) primarily capture distributed prediction error signals that are particular to each orientation (e.g., Bastos et al., 2012), then reduced classification performance for expected stimuli could be interpreted as reflecting suppressed prediction error signalling (e.g., Richter et al., 2022). Conversely, higher performance for expected stimuli would imply that such error signals are *enhanced* relative to other sources of stimulus-evoked neural activity (e.g., Kok et al., 2012; Yon et al., 2018). If one instead assumes that the stimulus-evoked responses used to train classifiers predominantly reflect the activity of prediction (also called representation) units, then higher classification accuracy would instead suggest relatively larger prediction unit responses. Alternatively, increases in classification performance may specifically arise when an observer’s expectations are violated, which would be consistent with a reactive orienting response to better sample from novel or unusual events in one’s environment (Press et al., 2020; Alink & Blank, 2021).

Our finding of higher classification accuracy for the surprising (10% probability) compared to the expected (90% probability) condition provides some support for reactive orienting responses, particularly as the (statistically significant) effects occurred over a relatively late time window (280-300ms) as compared to the bulk of the afferent visual evoked response. One possibility is that there is a threshold of surprisal that is required to trigger an orienting response (as described in Press et al., 2020). Our findings remain to be replicated (using a similar or wider range of appearance probability conditions) before strong support for this type of thresholded process can be inferred.

### 4.3. Stimulus Repetition Effects

In contrast to the apparent absence of expectation effects, we observed clear within-trial stimulus repetition effects on grating-evoked ERPs. This included an early (−75ms) onset effect (also reported in Tang et al., 2018) and a slightly later (−150-170ms) opposite polarity effect over occipital electrodes. Stimulus repetition effects could also be detected using MVPA over the same time windows. These findings are another example of clear adaptation (i.e., stimulus repetition) effects in the absence of detectable stimulus expectation effects (e.g., Kaliukhovich & Vogels, 2011, 2014; Solomon et al., 2021; den Ouden et al., 2023). However, when we trained classifiers to discriminate between S2 grating orientations using data from the randomised presentation blocks, we did not observe differences in classification performance between repeated and alternating conditions. Our results provide further support for the notion that adaptation and expectation-related phenomena are distinct and separable, contrary to predictive coding-based explanations of adaptation in the visual system (e.g., Summerfield et al., 2008; Auksztulewicz & Friston, 2016; discussed in Feuerriegel, 2024). We also demonstrate that repetition effects as detected using scalp EEG have a short onset latency, which may reflect response differences in orientation-selective neurons within early visual cortex (e.g., Patterson et al., 2014).

Our finding of similar grating orientation classification accuracy for repeated and alternating S2 stimuli is consistent with Tang et al. (2018), who also did not observe stimulus repetition effects on classification performance measures. These results are not consistent with hypothesised sharpening of population responses (Desimone, 1996; Grill-Spector et al., 2006; Rideaux et al., 2023), nor are they consistent with contemporary models of adaptation that specify reductions in the activity of stimulus-selective excitatory neurons that are tuned to the repeated grating orientation (e.g., Solomon & Kohn, 2014; Vogels, 2016; Whitmire & Stanley, 2016). It is somewhat puzzling why stimulus repetition leads to clear effects on ERP amplitudes, but small or absent effects on grating orientation-selective patterns as measured using scalp EEG. Please note that, in the current study, the repeated and alternating S2 gratings differed systematically with respect to the orientation of the preceding grating (which was the same or different, respectively). It is possible that late-emerging patterns evoked by the S1 gratings may have influenced classification in the period following S2 grating onset in this design. Although we did not observe clear indications of above-chance decoding beyond −400ms post S1 grating onset (Figure 6A, see also Grootswagers et al., 2024), late-arising effects cannot be ruled out entirely.

We also caution that longer latency repetition effects on ERPs (between −350-600ms) appeared to have posterior locus over visual cortex but may be (at least partly) due to faster RTs for repetition as compared to alternation trials. This typically co-occurs with different latencies of parietal ERP components (such as the centro-parietal positivity, O’Connell et al., 2012) that are time-locked to the decision and corresponding motor response (for example effects on ERPs see Sun et al., 2024). This is also important to consider in probabilistic cueing designs where RTs systematically differ across expected and surprising stimuli.

In contrast to den Ouden et al. (2023) and other work (e.g., Kaliukhovich & Vogels, 2014; Peter et al., 2021) that presented complex stimuli such as faces, animals, and objects, we did not find strong evidence of across-trial repetition effects for cues or gratings. Although long-lasting stimulation history effects have been observed for drifting gratings (Fritsche et al., 2022), it appears that such effects on EEG signals are more difficult to detect when brief, static gratings are presented. However, we note that it is still important to avoid stimulation history confounds that may arise from across-trial repetition effects.

### 4.4. Limitations

There are several caveats that should be considered when interpreting our results. First, the scope of the evidence reviewed here is limited to the visual system. Expectation effects have been reported in auditory probabilistic cueing designs (e.g., Todorovic & de Lange, 2012) and BOLD signals in the insula and other areas outside the visual system have been reported to covary with quantitative metrics of uncertainty and surprise (Preuschoff et al., 2008; Mohr et al., 2010; Loued-Khenissi et al., 2020). In addition, the EEG signals measured in our study are not sufficiently sensitive to differences in electrical activity within deep brain structures such as the striatum. The lack of EEG effects here does not imply that these areas are insensitive to manipulations of stimulus appearance probability.

In addition, participants in our study learned the relevant cue-stimulus associations *before* they commenced the probabilistic cueing task. This contrasts with other studies in which participants learned cue-stimulus associations via exposure across the course of the experiment (e.g., Summerfield et al., 2011; Hall et al., 2018). Our experiment was designed to test for effects associated with the consequences of already learned expectations, rather than neural correlates of belief formation and updating that might contribute to expectation effects in other work. Under some formulations (e.g., Kwisthout et al., 2017) the precision weighting of a prediction error will be low when stimulus appearance probabilities are fully known in advance. Although we tested for neural correlates of prediction error *magnitude*, our manipulations may not have produced large variations in precision weighted prediction error across conditions (in contrast to auditory studies by Lieder et al., 2013; Garrido et al., 2016; LeCaignard et al., 2022). However, we note that other studies, in which stimulus probabilities were not known in advance, also did not provide evidence for stimulus appearance probability effects (e.g., Kaliukhovich & Vogels, 2011; Kok et al., 2017; Vinken et al., 2018; Solomon et al., 2021).

We also note that this study was focused on effects of cue-stimulus association learning, but it is possible that participants were not consistently relying on the cues to perform the task in the probabilistic cueing experiment. In our experiment the S1 gratings were directly relevant to the grating matching task. However, participants could have plausibly performed the task without relying heavily on the predictive cue-stimulus relationships (e.g., by focusing on the identity of the S1 gratings and ignoring the cues). Here, we note that it is highly unlikely that participants were closing their eyes during the presentation of the cues. We observed very clear cue-evoked ERPs that resemble typical waveforms following visual stimuli (as visible in Figure 4C, Supplementary Figures S2B and S3). We also monitored eyeblinks during EEG recording and, following standard practice, instructed participants to blink during the inter-trial interval. Importantly, we investigated whether learned probabilistic cue-stimulus associations (without additional task-related factors) were sufficient to produce differences in grating-evoked ERPs (consistent with similar research questions in prior work, e.g., Summerfield et al., 2008; Egner et al., 2010; Kaliukhovich & Vogels, 2011; Song et al., 2024). Attending to the cue stimuli could also have been beneficial for performing the task, as the cue onsets provided information about the timing of the subsequent, briefly presented S1 grating stimuli.

However, it is possible that probabilistic cueing effects may be specific to when participants rely on predictive cues to perform a task (e.g., Larsson & Smith, 2012; Richter & de Lange, 2019; Alink & Blank, 2021; but see Kok et al., 2017; Rungratsameetaweeana et al., 2018; Vinken et al., 2018). To our understanding, dominant predictive coding-based accounts of probabilistic cueing effects do not specify that effects should only be observed under those conditions. Theoretical models could be developed to more explicitly define concepts such as attention and task relevance, and to specify the conditions under which expectation effects should (and should not) be observed.

Here, we have assumed that the relevant predictions in our experiment relate to subjective appearance probabilities of specific grating orientations. If one instead assumes a higher-level prediction, such as ‘either a vertical or horizontal stimulus will appear’, then all S1 stimuli would be reclassed as expected stimuli (see Kwisthout et al., 2017). When testing for effects of predictions, one encounters a problem of multiple indeterminacy. Assumptions must be made about i.) which predictions are formed in a given context, ii.) how each prediction is weighted relative to other co-occurring predictions, and iii.) the linking functions between each type of prediction error and neural response measures. Consequently, our findings may be interpreted differently by those assuming different mixtures of predictions that are formed within our experiment.

We also used a somewhat restricted range of stimulus appearance probabilities (ranging from 10-90%). While this spans the range of stimulus appearance probabilities included in previous work, it is possible that effects of surprise may occur for stimuli that have even lower appearance probabilities. As some contemporary predictive processing models (e.g., Press et al., 2020) do not specify a fixed quantitative degree of expectation violation that is required for surprise-related responses to occur, future work could include highly unlikely stimuli to test for such responses while controlling for known confounds such as stimulus novelty (discussed in Feuerriegel et al., 2021).

We also included only two stimulus identities that could appear after each cue, meaning that participants could form expectations about both the expected and surprising stimulus identity following a given cue. This was done to avoid novelty-related confounds and did not prevent us from assessing effects of *relative* subjective appearance probability (relating to the hypothesis specified in Walsh et al., 2020). It is possible that expectation suppression effects may be observed when a large range of unexpected stimuli could plausibly appear (discussed in Rostalski et al., 2020; Feuerriegel et al., 2021), however, to our knowledge this type of stimulus predictability does not appear to substantively modulate visual stimulus-evoked ERPs when novelty is equated across conditions (den Ouden et al., 2023).

### 4.5. Conclusion

The absence of detectable expectation effects observed in our study, as investigated using both univariate and multivariate analysis techniques, does not support the hypothesis that responses of stimulus-selective neurons in the visual system inversely scale with subjective appearance probability (Walsh et al., 2020). This does not support variants of predictive coding models that specify expectation suppression effects (e.g., Friston, 2005, 2010; Clark, 2013; Summerfield & de Lange, 2014). Our findings also do not support proposals that distributed responses across visual cortex (commonly termed stimulus representations) are modified by expectations about the probability of stimulus appearance (Kok et al., 2012; Richter et al., 2022).

## Acknowledgements

This work was supported by an Australian Research Council Discovery Early Career Researcher Award to D.F. (ARC DE220101508). Funding sources had no role in study design, data collection, analysis or interpretation of results. We thank Lara Aguila for insightful discussions relating to experiment design.

