## Supplementary Material for "Limited evidence for probabilistic cueing effects on grating-evoked event-related potentials and orientation decoding performance"

### 1. Event-Related Potential Results for the Parieto-Occipital Region of Interest

At parieto-occipital electrodes (PO7/8, P7/8/9/10), we did not find any statistically significant differences in ERP amplitudes across any of the compared stimulus appearance probability conditions after correcting for multiple comparisons. Supplementary Figure S1 displays the group average ERPs for each set of compared conditions, the difference waves, standardised Cohen's d effect size estimates, and Bayes factors in favour of the alternative hypothesis. Although there were brief periods over which Bayes factors rose above 1 (generally corresponding to $p < .05$ when using the defined Cauchy prior distribution) for comparisons between surprising (10% probability) and expected (90%) and neutral (50%) conditions (Figures S1A and S1E), these did not survive correction for multiple tests in our frequentist analyses. Bayes factors were generally less than 1 (indicating preferential support for the null hypothesis) for other comparisons (Figures S1B-D and S1F).

At parieto-occipital channels we observed within-trial grating image repetition effects within three distinct time windows, spanning 74-104ms (cluster $p = .031$), 336-486ms (cluster $p < .001$), and 518-600ms (cluster $p = .004$) from S2 grating onset. Group-averaged ERPs for repeated and alternating S2 gratings are shown in Supplementary Figure S2A. We did not observe any effects of cue image repetition across trials (Figure S2B). However, we did observe ERP differences between S1 gratings that were preceded by S2 gratings in the previous trial of the same as compared to different orientations, over time windows spanning 191-225ms (cluster $p$ = .049) and 438-469ms (cluster $p = .042$, Figure S2C).

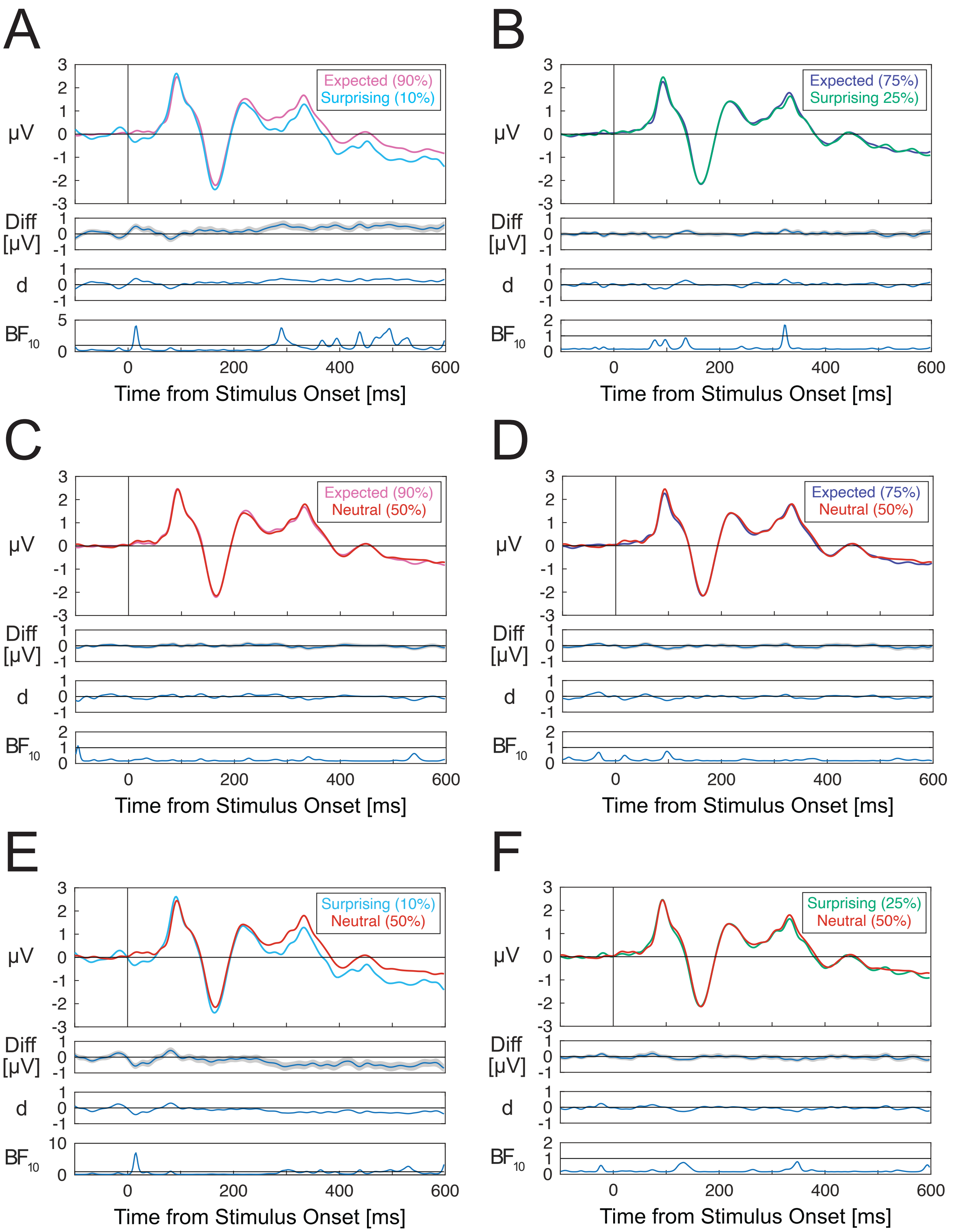
A
B
C
D
E
F
μV
Diff [μV]
d
$BF_{10}$
Time from Stimulus Onset [ms]
Expected (90%)
Surprising (10%)
Expected (75%)
Surprising 25%)
Neutral (50%)
Surprising (25%)

**Supplementary Figure S1.** Group-averaged ERPs evoked by S1 gratings within different stimulus appearance probability conditions. ERPs are averaged across channels within the parieto-occipital region of interest including channels PO7/8, P7/8 and P9/10. A) Expected (90%) – surprising (10%) ERP amplitude differences. B) Expected (90%) – surprising (25%) differences. C) Expected (90%) – neutral (50%) differences. D) Expected (90%) – neutral (25%) differences. E) Neutral (50%) – surprising (10%) differences. F) Neutral (50%) – surprising (25%) differences. ERPs for each set of compared conditions are displayed along with difference waves (with shading denoting standard errors), Cohen's d effect size estimates, and Bayes factors in favour of the alternative hypothesis. Horizontal lines in Bayes factor plots denote $BF_{10} = 1$, indicating a lack of preferential support for either the alternative or null hypothesis.

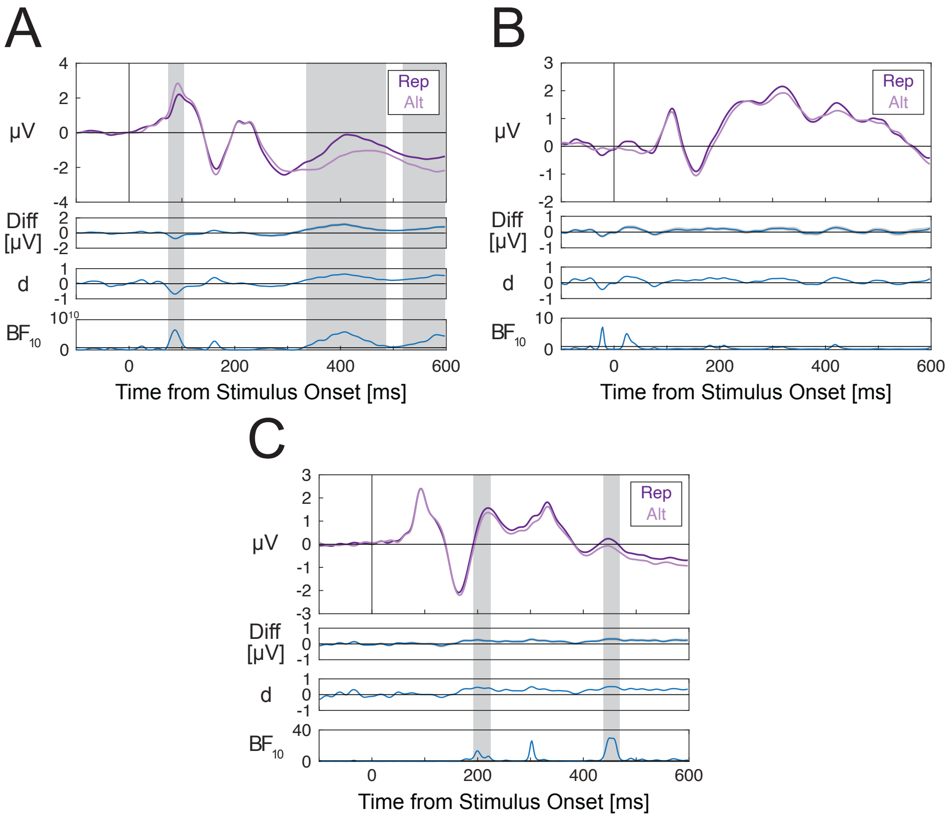


**Supplementary Figure S2**. Within- and across-trial repetition effects for S2 gratings, cues, and S1 gratings. ERPs are averaged across channels within the parieto-occipital region of interest including channels PO7/8, P7/8 and P9/10. A) S2 grating-evoked ERPs depending on whether the S2 grating was the same orientation as the preceding S1 grating in the same trial (repetition/Rep) or a different orientation (alternation/Alt). B) Cue-evoked ERPs depending on whether the cue image was the same as the cue in the previous trial (Rep) or a different cue image (Alt). C) S1 grating-evoked ERPs depending on whether the S2 grating in the previous trial was the same (Rep) or a different orientation (Alt). For each pair of compared conditions, ERPs are displayed along with difference waves (shading denoting standard errors), Cohen's d effect size estimates, and Bayes factors in favour of the alternative hypothesis. Note that for A) Bayes factors are plotted on logarithmic scales due to the wide ranges of values across the time-course of the stimulus-evoked response. Horizontal lines in Bayes factor plots denote $BF_{10} = 1$, indicating a lack of preferential support for either the alternative or null hypothesis. Grey shaded areas denote time windows of statistically significant differences after correcting for multiple comparisons.

### 2. Comparisons of ERPs Evoked by Different Cue Types

At occipital electrodes (Oz/O1/O2/POz/Iz), we did not find any statistically significant differences in ERP amplitudes across any of the compared cues after correcting for multiple comparisons. Supplementary Figure S3 displays the group average ERPs for each set of compared conditions, the difference waves, standardised Cohen's d effect size estimates, and Bayes factors in favour of the alternative hypothesis. There were no apparent differences between 50% and 90% grating appearance probability cues (Figure S3A), 50% and 75% cues (Figure S3B), or 75% and 90% cues (Figure S3C). Although there were brief periods over which Bayes factors rose above 1 (generally corresponding to $p < .05$ when using the defined Cauchy prior distribution), these did not survive correction for multiple tests in our frequentist analyses. Bayes factors were generally less than 1 (indicating preferential support for the null hypothesis).

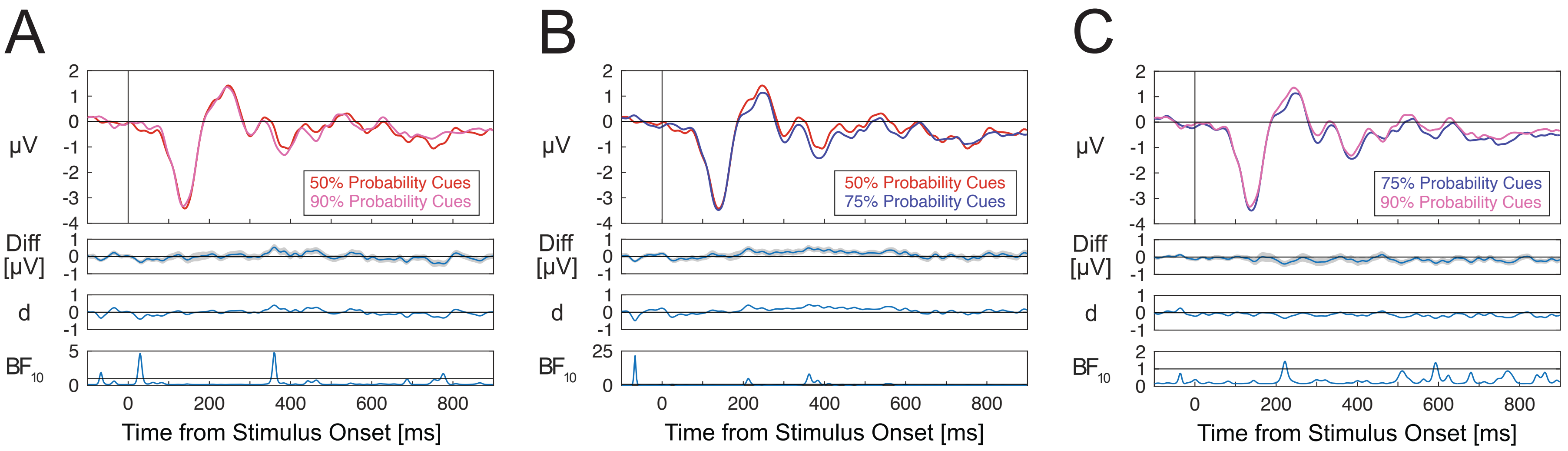


**Supplementary Figure S3**: ERPs evoked by cues that signalled different grating appearance probability conditions. A) ERPs evoked by cues that signalled either a 50% or 90% probability grating orientation. B) ERPs evoked by cues that signalled either a 50% or 75% probability grating. C) ERPs evoked by cues that signalled either a 75% or 90% probability grating. ERPs are averaged across channels within the occipital ROI including channels Oz/1/2, POz and Iz. ERPs for each set of compared conditions are displayed along with difference waves (with shading denoting standard errors), Cohen's d estimates and Bayes factors. Horizontal lines in Bayes factor plots denote $BF_{10}$ = 1, indicating a lack of preferential support for either the alternative or null hypothesis.

### 3. Classification of Grating Orientation - Comparisons Across S1 Grating Appearance Probability Conditions Using Pre-Cue Baselines

We also performed S1 grating orientation classification analyses using epochs that were baseline-corrected to the 100ms pre-cue time window (rather than a pre-S1 grating time window). When averaging across S1 appearance probability conditions we observed above-chance decoding performance (after correction for multiple comparisons) over two narrow time windows spanning 216-235ms and 300-318ms (Supplementary Figure S4). We did not observe any differences in classification performance across S1 grating probability conditions (Supplementary Figure S5).

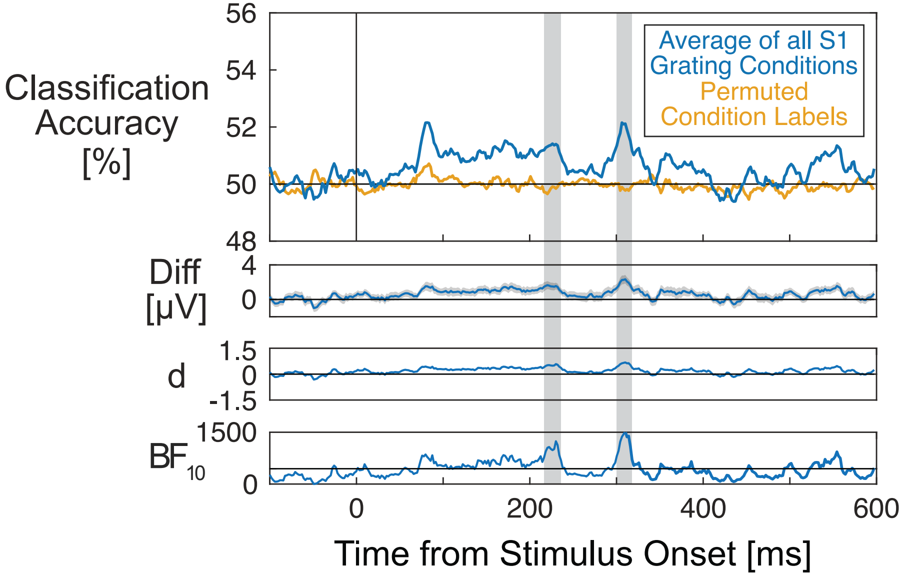

**Supplementary Figure S4.** Classification performance results for classifiers trained to discriminate between S1 grating orientations for epochs that were baseline-corrected using the pre-cue interval. Classification accuracy and permuted-labels classification accuracy is displayed along with difference waves (shading denoting standard errors), Cohen's d effect size estimates, and Bayes factors in favour of the alternative hypothesis. Grey shaded areas denote time windows of statistically significant differences after correcting for multiple comparisons. Horizontal lines in Bayes factor plots denote $BF_{10} = 1$, indicating a lack of preferential support for either the alternative or null hypothesis.

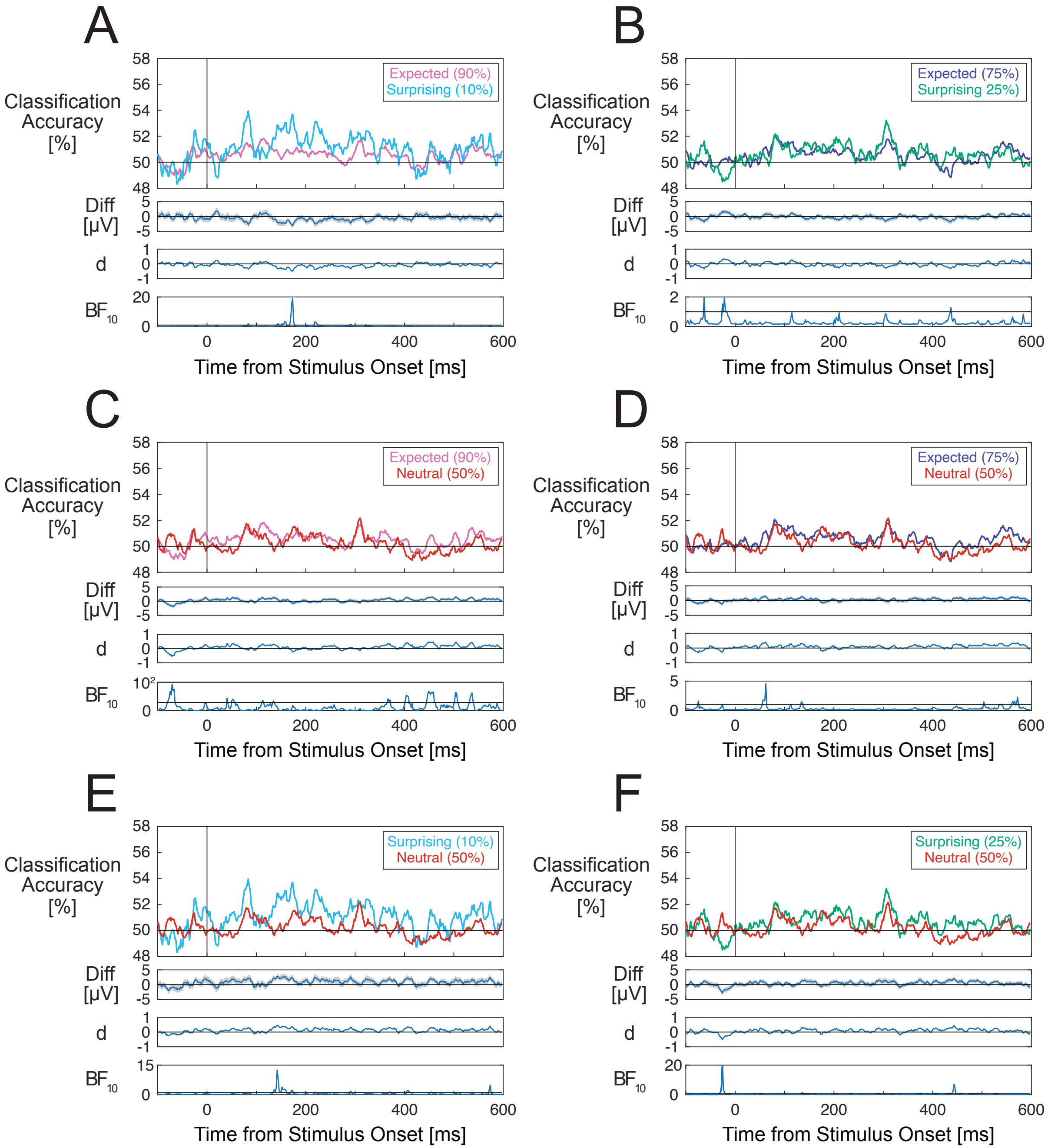
A
Expected (90%)
Surprising (10%)
Classification Accuracy [%]
Diff [μV]
d
BF10
Time from Stimulus Onset [ms]
B
Expected (75%)
Surprising 25%)
C
Expected (90%)
Neutral (50%)
D
Expected (75%)
Neutral (50%)
E
Surprising (10%)
Neutral (50%)
F
Surprising (25%)
Neutral (50%)

**Supplementary Figure S5.** Classification performance results for classifiers trained to discriminate between S1 grating orientations for epochs that were baseline-corrected to the pre-cue interval. A) Classification performance for expected (90%) and surprising (10%) S1 grating conditions. B) Classification performance for expected (75%) and surprising (25%) conditions. C) Classification performance for expected (90%) and neutral (50%) conditions. D) Classification performance for expected (75%) and neutral (50%) conditions. E) Classification performance for surprising (10%) and neutral (50%) conditions. F) Classification performance for surprising (25%) and neutral (50%) conditions. For each pair of conditions, classification accuracy is displayed along with difference waves (shading denoting standard errors), Cohen's d effect size estimates, and Bayes factors in favour of the alternative hypothesis. Note that for C) Bayes factors are plotted on a logarithmic scale due to the wide ranges of values across the time-course of the stimulus-evoked response. Horizontal lines in Bayes factor plots denote $BF_{10} = 1$, indicating a lack of preferential support for either the alternative or null hypothesis.

### 4. Temporal Generalisation Analyses for Classification of Grating Orientations

We also performed exploratory temporal generalisation analyses. Classifiers trained at a given time point relative to grating stimulus onset were assessed on their ability to classify grating orientations using data at all other time points.

Classifiers trained using vertical and horizontal grating-evoked single-trial ERPs from the randomised presentation blocks were used to discriminate between horizontal and vertical S1 and S2 gratings. Numbers of trials per grating orientation were equated for the classifier training datasets. In cases of an imbalanced number of trials across grating orientations in the randomised presentation blocks, a random subset of trials was removed from the condition with more trials. To maximise the amount of available testing data, trial balancing was not done for the S1 and S2 gratings. However, this should in principle not bias classifier accuracy as the training dataset was balanced. To account for any general differences in ERPs across randomised blocks and the probabilistic cueing experiment, the trial-average ERP was subtracted from each trial in the training dataset. ERPs were averaged within each grating orientation condition for the S1 and S2 grating epochs used to test the classifiers, and the average of these was subtracted from each single trial of EEG data used to test the classifiers.

We tested for above-chance classification accuracy at the group level using one-sample t-tests (two-tailed, alpha = 0.01). In this case, two-tailed tests were used as below-chance classification accuracy is possible when training and testing using data from different time points. We tested for differences in accuracy across S1 appearance probability and S2 repetition/alternation conditions using paired-samples t-tests (two-tailed, alpha = 0.01). Tests were performed for each training/test time point combination (total of 128,164 tests for each temporal generalisation analysis). As these analyses were exploratory and designed to be sensitive enough to identify candidate effects for future replication, we do not encourage strong inferences to be drawn based on these results. There is also likely to be at least some proportion of false-positive results in this data.

Classification accuracy for the average of the S1 probability conditions is shown in Supplementary Figure S6. A contiguous region of above-chance decoding was observed that was limited to the diagonal (i.e., regions in which the training and testing time points were close in time). This time-course was similar to the results of the analyses using the same training and testing time points for classification depicted in Figure 6A of the corresponding paper.

Comparisons across S1 appearance probability conditions (depicted in Supplementary Figure S7A-F) yielded only sparse, scattered patterns of training/testing time point combinations at which classification accuracy was significantly different using the set alpha level. These do not appear to be genuine effects, which tend to extend over longer periods of time due to the substantial degree of autocorrelation in EEG data. We also did not find clear evidence of classification accuracy differences across repeated and alternating S2 gratings (Supplementary Figure S8).

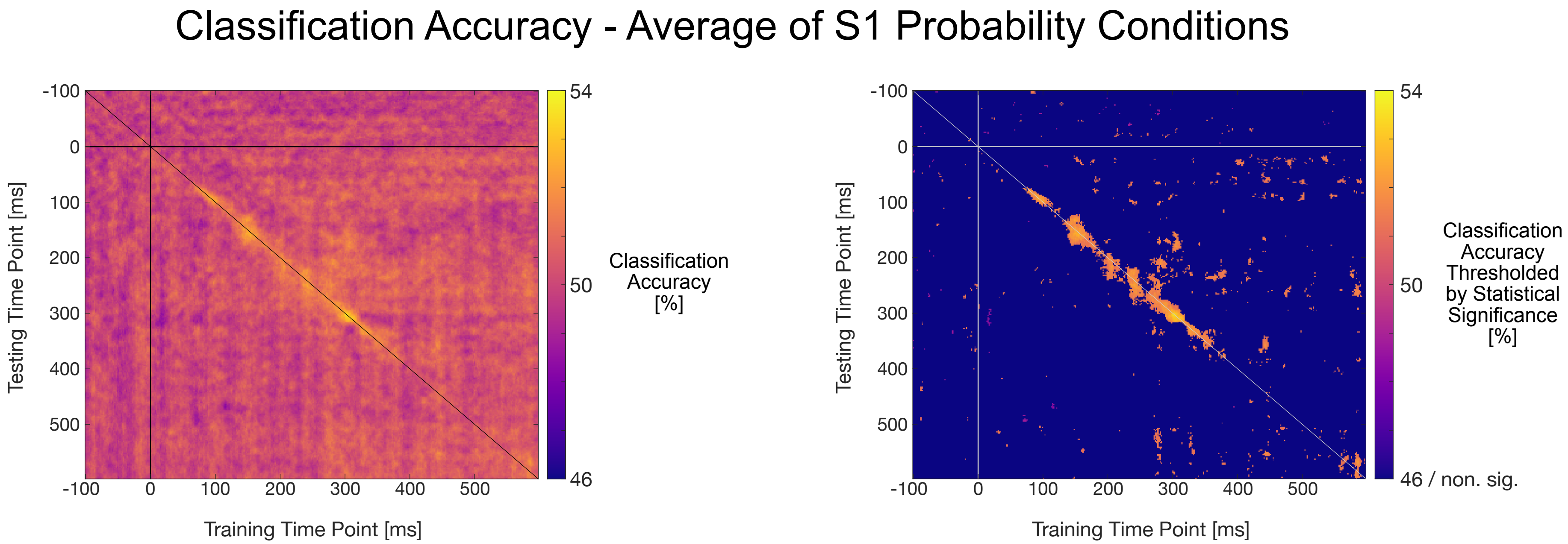


**Supplementary Figure S6.** Classification performance for classifiers trained to discriminate between S1 grating orientations. Classifiers were trained on data from the randomised presentation blocks. Temporal generalisation plots show performance for each training/testing time point combination relative to grating stimulus onset. The left plot shows group mean classification accuracy. The right plot shows group mean classification accuracy thresholded by statistical significance ($p < .01$, uncorrected).

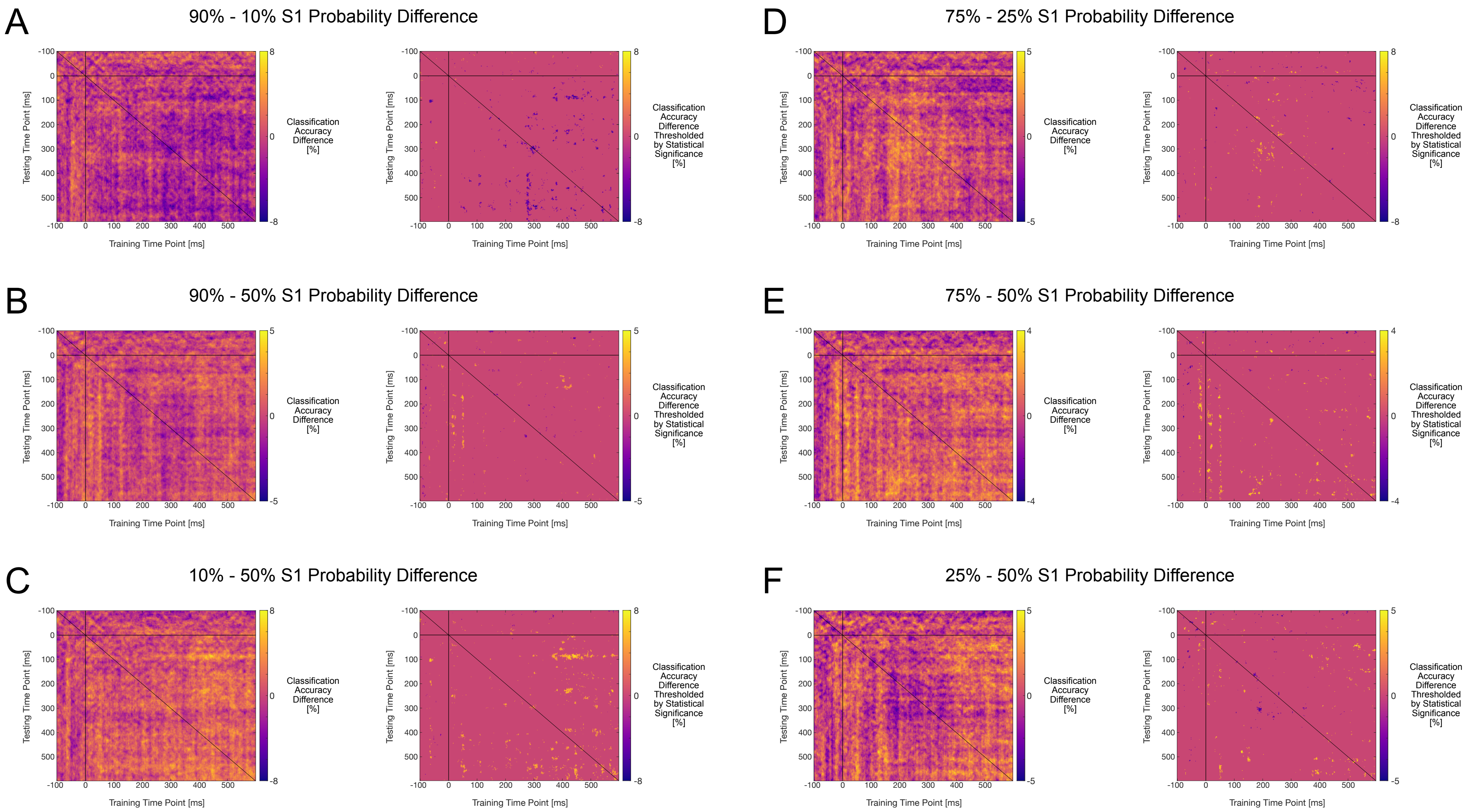
A
90% - 10% S1 Probability Difference
B
90% - 50% S1 Probability Difference
C
10% - 50% S1 Probability Difference
D
75% - 25% S1 Probability Difference
E
75% - 50% S1 Probability Difference
F
25% - 50% S1 Probability Difference
Testing Time Point [ms]
Training Time Point [ms]
Classification Accuracy Difference [%]
Classification Accuracy Difference Thresholded by Statistical Significance [%]

**Supplementary Figure S7.** Classification performance differences across S1 appearance probability conditions, for classifiers trained to discriminate between S1 grating orientations. Classifiers were trained on data from the randomised presentation blocks. Temporal generalisation plots show performance for each training/testing time point combination relative to grating stimulus onset. A) 90% vs. 10% condition differences. B) 90% vs. 50% condition differences. C) 10% vs. 50% condition differences. D) 75% vs. 25% condition differences. E) 75% vs. 50% condition differences. F) 25% vs. 50% condition differences. The left plots show group mean classification accuracy. The right plots show group mean classification accuracy thresholded by statistical significance ($p < .01$, uncorrected).

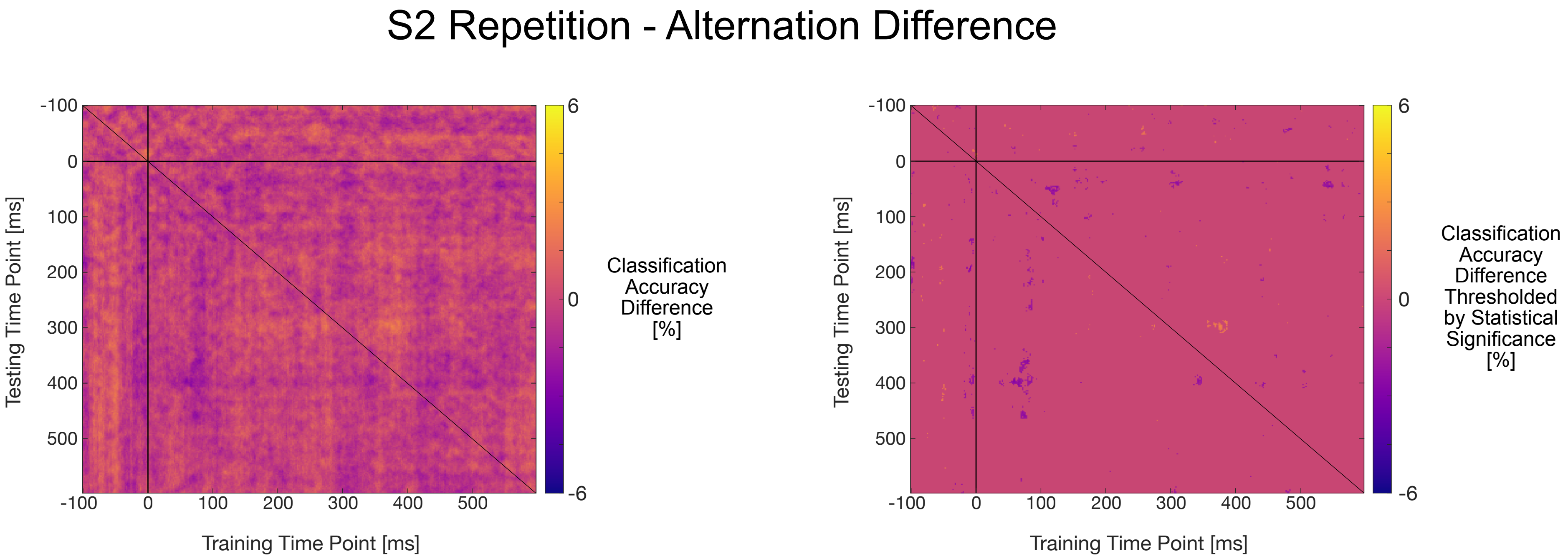


**Supplementary Figure S8.** Classification performance differences across S2 repetition/alternation conditions, for classifiers trained to discriminate between S2 grating orientations. Classifiers were trained on data from the randomised presentation blocks. Temporal generalisation plots show performance for each training/testing time point combination relative to grating stimulus onset. The left plot shows group mean classification accuracy. The right plot shows group mean classification accuracy thresholded by statistical significance ($p < .01$, uncorrected).
